# Unraveling the Phenotypic States of Human innate-like T Cells: Comparative Insights with Conventional T Cells and Mouse Models

**DOI:** 10.1101/2023.12.07.570707

**Authors:** Liyen Loh, Salomé Carcy, Harsha S. Krovi, Joanne Domenico, Andrea Spengler, Yong Lin, Joshua Torres, William Palmer, Paul J. Norman, Matthew Stone, Tonya Brunetti, Hannah V. Meyer, Laurent Gapin

**Author notes:** Corresponding author: Dr. Laurent Gapin, Department of Immunology and Microbiology, University of Colorado Anschutz Medical campus, 12800 E. 19th Ave., Aurora, CO, 80045, USA. Dr. Hannah Meyer, Simons Center for Quantitative Biology, Cold Spring Harbor Laboratory, Cold Spring Harbor, NY, USA., *E-mail addresses. These authors contributed equally (Liyen Loh, Salomé Carcy, Hannah V. Meyer, Laurent Gapin).

## Abstract

The “innate-like” T cell compartment, known as T_inn_, represents a diverse group of T cells that straddle the boundary between innate and adaptive immunity, having the ability to mount rapid responses following activation. In mice, this ability is acquired during thymic development. We explored the transcriptional landscape of T_inn_ compared to conventional T cells (T_conv_) in the human thymus and blood using single cell RNA sequencing and flow cytometry. We reveal that in human blood, the majority of T_inn_ cells, including iNKT, MAIT, and Vδ2^+^Vγ9^+^ T cells, share an effector program characterized by the expression of unique chemokine and cytokine receptors, and cytotoxic molecules. This program is driven by specific transcription factors, distinct from those governing T_conv_ cells. Conversely, only a fraction of thymic T_inn_ cells displays an effector phenotype, while others share transcriptional features with developing T_conv_ cells, indicating potential divergent developmental pathways. Unlike the mouse, human T_inn_ cells do not differentiate into multiple effector subsets but develop a mixed type I/type III effector potential. To conduct a comprehensive cross-species analysis, we constructed a murine T_inn_ developmental atlas and uncovered additional species-specific distinctions, including the absence of type II T_inn_ cells in humans, which implies distinct immune regulatory mechanisms across species. The study provides insights into the development and functionality of T_inn_ cells, emphasizing their role in immune responses and their potential as targets for therapeutic interventions.

## Introduction

The immune system is a complex network that offers protection against pathogens through two primary classifications: “innate” and “adaptive” immunity. Innate immunity involves pre-established reactions driven by fixed, germline-encoded immune receptors, while adaptive immunity relies on the rearrangement and alteration of germline DNA to produce unique T and B cell antigen receptors which detect molecules derived from pathogens.

Conventional CD4^+^ and CD8^+^ T cells (T_conv_) play a crucial role in the adaptive immune response. They express T cell antigen receptors (TCRs) that recognize linear peptide fragments presented by major histocompatibility complex class I or II (HLA class I or HLA class II) proteins. Upon encountering their cognate antigens, these T cells undergo significant transcriptional and epigenetic changes, leading to the secretion of pro-inflammatory cytokines, chemokines and acquisition of cytotoxic capability that promote pathogen clearance. This process results in the formation of memory T cells, which are primed to respond rapidly upon reencountering the pathogen. Thus, T_conv_ cells within the circulation are heterogeneous and surface markers such as CCR7, CD45RA, and CD62L are commonly used to classify them into naïve (T_n_), central memory (T_cm_), effector memory (T_em_), and terminally differentiated effector memory (T_emra_) subsets (Jameson and Masopust, 2018; Kaech and Cui, 2012; Sallusto et al., 1999).

Recent studies have challenged the idea that somatic recombination is exclusively linked to adaptive immunity. Over the last 20 years, T-cell populations with TCRs that remain consistent among individuals and develop effector functions without prior pathogen exposure were discovered (Godfrey et al., 2015; Mayassi et al., 2021). These “innate-like” T-cell populations (T_inn_), such as invariant natural killer T (iNKT) cells, mucosal-associated invariant T (MAIT) cells, and γδ T cells, account for a significant portion of human T cells, estimated to be between 10 and 20% (Godfrey *et al*., 2015). They serve vital roles in host defense and immune homeostasis (Chandra and Kronenberg, 2015; Godfrey et al., 2019; Hayday, 2019).

T_inn_ cells originate from the same thymic progenitor cells as adaptive T cells but possess several distinguishing features that set them apart from T_conv_ cells. Firstly, they do not recognize peptides presented by HLA class I or class II. iNKT cells express semi-invariant αβ TCRs characterized in humans by a TRAV10-TRAJ18 Vα chain coupled with a limited Vβ repertoire (TRVB25) and recognize self- and foreign-lipid antigens presented by the non-polymorphic HLA-like molecule, CD1D (Matsuda et al., 2008). They are specifically detected using CD1D tetramers loaded with the cognate lipid antigen α-galactosylceramide (αGC) (Benlagha et al., 2000; Matsuda et al., 2000). MAIT cells are similarly characterized through the usage of a semi-invariant TCR α chain associating TRAV1-2 with TRAJ33 (or TRAJ20, or TRAJ12) that is paired with a limited number of TRBV chains (Legoux et al., 2017). The TCRs formed by these combinations can be detected with tetramers of the MAIT restricting molecule, MR1, when loaded with 5-(2-oxopropylideneamino)-6-d-ribitylaminouracil (5-OP-RU), a derivative of the microbial vitamin B2 precursor 5-Amino-6-(D-ribitylamino)uracil (5-A-RU) (Reantragoon et al., 2013). γδ T cells express TCRs encoded by TRGV and TRDV gene segments but the specific antigen-presenting elements responsible for their development or activation remain unknown. A major γδ T-cell population bearing Vδ2-Vγ9 TCRs is activated by self- and foreign phosphoantigens in conjunction with transmembrane butyrophilin-family receptors BTN2A1-BTN3A1-BTN3A2 complex (Harly et al., 2012; Karunakaran et al., 2020; Rigau et al., 2020). The antigens recognized by other human γδ T-cell populations remain unclear (Deseke and Prinz, 2020). In summary, T_inn_ expand the spectrum of antigens detectable by T cells, enhancing the immune system’s ability to recognize and respond to a diverse array of threats.

The conservation of T_inn_ cells throughout mammalian evolution indicates a crucial and nonredundant role for these subsets in the immune system (Harly et al., 2022). This importance may be attributed to their innate characteristics displayed during inflammation and infection, such as rapid activation kinetics without prior pathogen exposure and the ability for antigen receptor-independent activation. Inflammatory cytokines, including IL-12, IL-18, and type I interferons, can activate T_inn_ cells even in the absence of simultaneous signaling through their TCRs (Leite-De-Moraes et al., 1999; Ussher et al., 2014).

In mice, the rapid effector capacity of T_inn_ cells is due to a unique transcriptional program formed during their development in the thymus, distinguishing them from conventional T cells (Baranek et al., 2022; Krovi et al., 2022). Analogous to CD4 T_conv_ cells, which can be polarized by cytokines into T helper (Th) phenotypes such as Th1, Th2, and Th17 that secrete IFNγ, IL-4, and IL-17 respectively, mouse T_inn_ cells diverge into distinct, terminally differentiated, subsets that can be readily identified based on the expression of specific transcription factors like PLZF, GATA3, T-bet, and RORγt (Lee et al., 2013). Additionally, mouse iNKT subsets produce cytokines at steady state, directly affecting surrounding cells in the microenvironment and the development and polarization of T_conv_ cells (Breed et al., 2022; Cui et al., 2022; Lee *et al*., 2013). This implies that T_inn_ cells may function as gatekeepers, ensuring proper T cell development and maturation.

While studies in mice have delineated the developmental trajectories of T_inn_ and analyses of distinct subsets of peripheral human T_inn_ cells have shed light on the developmental stages of human Vδ2-Vγ9 (Perriman et al., 2023), and functional subtypes of human MAIT cells (Chandra et al., 2023; Garner et al., 2023), a comprehensive picture spanning development and peripheral function across T_inn_ and T_conv_ is lacking. In this study, we utilized the unbiased potential of single-cell genomics combined with flow cytometry to assess the range of phenotypic states T_conv_ and T_inn_ cells can adopt *in vivo* in the human thymus and blood. We uncovered that the majority of postnatal human thymic T_inn_ cells exhibit a transcriptome akin to that of naive CD4^+^ or CD8^+^ T_conv_ cells. Only a fraction of thymic T_inn_ cells show a transcriptional signature indicative of an “effector” state. Conversely, most adult blood T_inn_ cells display an effector transcriptome. While T_conv_ cells exhibit a continuum of transcriptional states, spanning from naive to central and effector memory T cells, T_inn_ cells express a distinct transcriptional program shared among iNKT, MAIT and Vδ2Vγ9 T cells. However, unlike the mouse, human T_inn_ cells do not differentiate into functionally distinct subsets; instead, they develop an effector program with mixed type 1/type 3 effector potential. Notably, our study demonstrates that the major transcription factors governing the human T_inn_ program are also expressed in mouse T_inn_ cells, although species-specific differences were also apparent. Finally, our study highlights differences in the pattern of CD1D expression in the thymus between the two species, which could potentially impact the maturation process of iNKT cells in humans.

## Results

### Single-Cell RNA Sequencing Analysis of T Cell Maturation and Post-Maturation Stages in Humans

To comprehensively explore the transcriptional profile of T_inn_ and T_conv_ throughout their maturation and post-maturation stages in humans, we conducted single-cell RNA sequencing on tetramer-sorted iNKT cells (PBS57-CD1d tetramer^+^, TRAV10^+^), MAIT (5-OP-RU-MR1 tetramer^+^, TRAV1-2^+^), and total γδ T cells, in addition to single positive (SP) CD4 and CD8 T_conv_ cells derived from four pediatric thymus and twelve adult blood samples. Tetramer-based sorting of iNKT and MAIT cells was used to enrich these rare cell populations and labeling by DNA-barcoded antibodies (“hashtags”) allowed us to confidently assign cell identities based on TCR specificity (Supplementary Fig 1). The sorted cell populations were then pooled across batches and loaded onto a BD Rhapsody cartridge, which allowed for single-cell capture and library construction. A subset of samples was also subjected to VDJ sequencing (Fig 1A, Supplementary Table I). A total of 78,607 cells (37,369 cells from pediatric thymus and 41,238 cells from adult blood) passed quality control (see methods) and were integrated into a combined reference dataset that minimized batch-associated variation while preserving tissue-specific differences (Fig. 1B, C and Supplementary Fig 2A, B). To identify and characterize subpopulation structures, we used unsupervised graph-based clustering, which led to the assignment of 18 distinct and stable clusters (Fig 1C), as assessed by repeated sampling and reassignment of the cells to clusters (Supplementary Fig 2C). Distinct niches were identified, primarily separating into thymus (clusters 0 through 9) or blood-associated regions (clusters 12 through 17), with a transitioning niche exhibiting an equal proportion of cells from both thymus and blood tissues (clusters 10 and 11) (Fig 1D & E). Cells in this space represent naïve T cells that are prepared to leave the thymus and/or just populated the blood, in agreement with their overrepresentation of an ‘egress’ gene signature (Sanchez Sanchez et al., 2022) (Fig 1F, Supplementary Table III).

**Figure 1:**
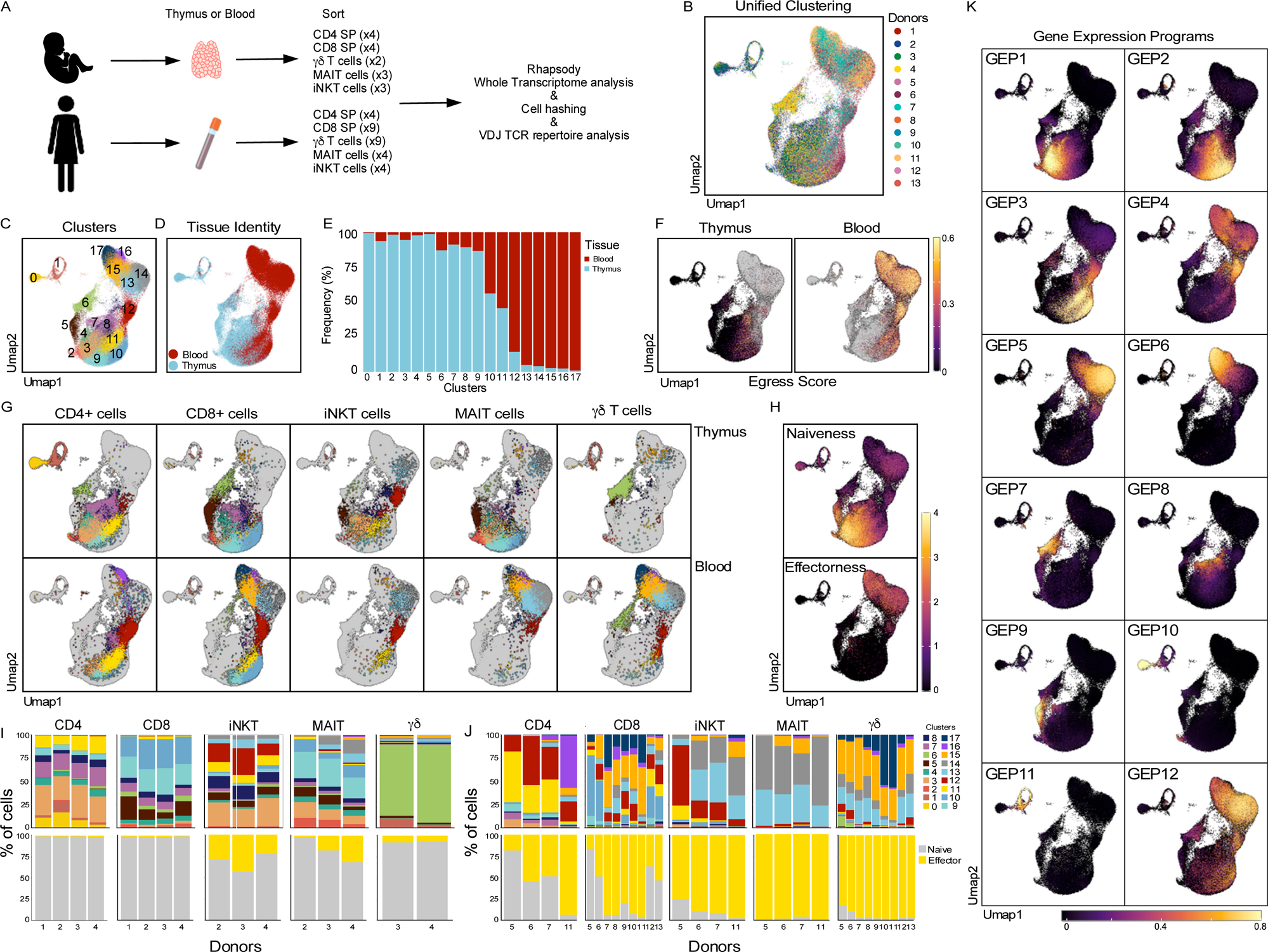
Integrative view on T_inn_ and T_conv_ development and peripheral function. A. Experimental set-up specifying donor type (postnatal/adult), tissue and sorted cell types. B. Harmony batch-corrected and integrated dataset across donors, tissues and cell types. C. Stable Louvain-derived cell clusters distributed across D. both blood and thymus-derived cells and E. their respective frequencies in these clusters. F. ‘Egress’ score on thymus and blood derived cells. G. Cells color-coded by cluster (as in C) and visualized by their hashtag-sorted cell type (columns) and the tissue they originated from (rows). H. Projection of naive and effector scores and the proportion of I. thymic and J. blood cell types per donor classified based on these scores (bottom row); top row shows analogous proportions by cell cluster (as in C). K. Gene expression programs (GEPs) in thymus and blood identified using cNMF. Sample numbers for all panels as depicted in A. Score defining genes as described in text.

The cell identities and/or transcriptional states of the clusters were determined using reference signature genes lists (Park et al., 2020), and the top five genes that characterize each cluster (Supplementary Fig 3A, B, Table II). Additionally, we used neighbor voting (Crow et al., 2018) with the cells from the human thymus atlas (Park *et al*., 2020) to assess the replicability of cell types and validate the assigned cell identities of the clusters that exhibited high similarity (Supplementary Fig 4). Starting at the beginning of T cell development, immature single positive (ISP) cells with a quiescent (cluster 0, from here on c0 and accordingly for other clusters) and cycling cell population (c1) were identified, along with double positive (CD4^+^CD8^+^, DP) thymocytes (c2). In humans, ISP cells, which precede the DP stage of development, express the CD4 molecule and were inadvertently sorted together with SP CD4 T cells. From DPs, thymocytes mature into CD4 single-positive (SP) and CD8 SP cells. Early stages of SP are characterized by the expression of the chemokine receptor *CCR9* (c3 for CD4 SP and c9 for CD8 SP), which is lost in later stages, concomitant with the expression of *CCR7* (c11 for CD4 SP, c10 for CD8 SP; Supplementary Fig 5A). Differential gene expression analysis comparing CD4 SP cells (c3 and c11) to CD8 SP cells (c9 and c10) further confirms these annotations, based on the overlap of the differentially expressed genes (DEGs) and previously defined signatures that distinguish CD4 SP from CD8 SP cells (Chopp et al., 2020) (Supplementary Fig 5B). Specialized lineages were detected in distinct regions in the two-dimensional UMAP space, including CD8αα cells with their distinct gene signature including expression of *GNG4* and *NUCB2*, thymic γδ T cells and regulatory T cells (T_regs_) with high expression of *FOXP3* (c5, c6 and c7, respectively). Other signaling states included cells with high levels of transcripts encoding transcription factors associated with TCR signaling (c4), such as *NR4A1*, *EGR1*, *EGR3*, and *NFKBID*, and were named “agonist”, cells with high expression of type I interferon signaling genes (*IFI6*, *MX1*, and *IFI44L* in c8), AP-1 transcription factors (*JUN*, *FOS*, *JUNB* in c12), and effector-encoding genes (*GZMK*, *GZMH*, *GZMB*, *PRF1*, and *CCL5*), suggesting involvement in effector functions of these cells (c13 through c17) were also found. Altogether, the clusters and their low-dimensional embedding displayed distinct transcriptional profiles, representing unique cell types (CD8αα, Tregs) as well as various stages of T cell development and maturation.

### Identification of the Gene Expression Programs that Characterize T cell populations in Thymus and Blood

Deciphering scRNA-seq data can be challenging due to the intricate nature of each cell’s gene expression pattern, which may encapsulate both its inherent identity and its present activity or role. To tackle this complexity, we applied consensus non-negative matrix factorization (cNMF) to project the high-dimensional data into lower-dimensional factors, enabling the identification of gene modules with similar biological functions that exhibit high correlations (Kotliar et al., 2019). We identified 12 distinct gene expression programs (GEPs; Fig 1K, Supplementary Table IV). To assess the contribution of each cell type to these GEPs, we separated each sample by cell type and tissue using the identifying tag (Fig 1G) and observed that some of these GEPs define activity programs shared across different cell types, while others are unique to specific cell clusters (Fig 1K). Specifically, GEPs 7 to 11 were associated with thymic γδ T cells, Tregs, thymic CD8αα T cells, quiescent ISP and proliferating ISP, respectively. We excluded GEP 12 from further analysis as it was driven by a batch effect (Supplementary Fig 6). GEP 1 and GEP 2, characterized by the presence of *CCR9* and *CCR7*, respectively, exhibited heightened activity in early and late developing thymic T cells. GEP3 was prominently expressed by naïve T cells. Among the remaining GEPs, three gene modules showed distinct distributions that overlapped with previously defined “effectorness” signatures, which are both exhibited in T_conv_ and T_inn_ (Fig 1G, H): GEP 4 showed high activity in cluster 12, GEP 5 exhibited high activity in clusters 13 and 14, while GEP 6 displayed the highest activity in clusters 15, 16, and 17 (Fig 1K). Leveraging insights from these gene modules, as detailed in the subsequent sections, we conducted an in-depth analysis of thymic and blood T cell populations, providing an integrated understanding of T cell differentiation and function.

### Unbiased Transcriptomic Analysis of Human T_inn_ Differentiation

To assess the distribution of each sorted T cell population across the 18 transcriptionally distinct clusters and GEPs, we separated each sample by cell type and tissue using the identifying tag (Fig 1G). Consistent with the gene signatures of each cluster, CD4^+^ thymic cells were predominantly found in clusters 0, 1, 2, 3, 4, 7, 8, and 11, while CD8^+^ thymic cells were primarily located in clusters 2, 5, 6, 7, 8, 9, and 10. Notably, the proportion of cells in each cluster was consistent across the four independent samples analyzed, with approximately 1% (1.1% ± 0.4%) of the cells populating clusters exhibiting an effector signature (Fig 1G, H & I). Unexpectedly, a substantial proportion of thymic iNKT cells were distributed across the same clusters as conventional CD4^+^ cells, while thymic MAIT cells predominantly shared clusters with conventional CD8^+^ cells (Fig 1G). Interestingly, thymic γδ T cells were transcriptionally distinct from all other cell types, with most of the cells occupying cluster 6 and GEP7 (Fig 1 G, K).

To delve further into the transcriptional heterogeneity of human thymic T_inn_ cells, we re-analyzed the iNKT and MAIT cell populations individually (Fig 2). We found seven stable clusters for both cell types (Fig 2A, E), and the proportion of cells in each cluster was consistent across donors (Fig 2B, F). We identified five major cell signatures that were shared across T_conv_, iNKT and MAIT cells (Supplementary Figure 3C, D). First, we observed a distinctive gene signature associated with CD8αα T cells (captured by GEP9, as illustrated in Fig 1K and Supplementary Fig 7, Tables V and VI). This signature was characterized by the heightened expression of genes including *NUCB2*, *MINDY2*, and *HIVEP3* (Supplementary Fig 3C, D), and intriguingly, it was observed in both iNKT and MAIT cells (termed NKT_c0 and MAIT_c1) (as shown in Fig 2C and G). Notably, this specific subset of thymic CD8^+^ iNKT cells, which also exhibited some PLZF expression while lacking *CD161*, *EOMES*, and *GZMK* expression (Fig 2M), could be readily identified using flow cytometry (Supplementary Fig 8). These findings imply the potential existence of a unique selection process for certain T_inn_ cells, possibly linked to their colonization of the gut epithelium. Second, we identified a similar pattern of expression for the *CCR9* and *CCR7* chemokine receptors, which serve as markers for early and late SP stages in T_conv_ cells, as well as in iNKT and MAIT cells (Fig 2M and Supplementary Fig 7). Initially, iNKT and MAIT cells exhibited an upregulation of *CCR9* in conjunction with *TOX* and *SATB1* (Supplementary Fig 3C, D), resembling the developmental program seen in early developing CD4 SP and CD8 SP cells, respectively. Subsequently, the elevated expression of *CCR7* marked cells that appeared to be at a more advanced developmental stage (termed NKT_c2 and MAIT_c4). These sequential waves of chemokine receptor expression align with gene modules GEP1 and GEP2 (Supplementary Fig 7), suggesting that they might be induced sequentially during the development of CD4, CD8, iNKT, and MAIT cells. These findings lend support to the notion that the human thymus harbors iNKT and MAIT cells with a transcriptome resembling that of developing naïve T_conv_ cells. The existence of such naïve populations (CD161^−^ EOMES^−^ GZMK^−^) of iNKT and MAIT cells in the human thymus were confirmed by flow cytometry (Supplementary Fig 8). Third, we discovered iNKT and MAIT cells characterized by upregulation of genes associated with type I interferon signaling such as *MX1* and *IFI6* (NKT_c3 and MAIT_c5, Supplementary Fig 3C, D), similar to CD4 and CD8 SP cells. Fourth, we detected SP-corresponding TCR signaling/AP-1 signatures. In iNKT cells, the upregulation of genes encoding AP-1 family transcription factors *FOS* and *JUN* correlated with expression of genes typically associated with this cell type, such as *ZBTB16* and *KLRB1*. These cells also expressed CD4 transcripts but not CD8A (NKT_c5, Supplementary Fig 3C). The TCR signature was more pronounced in MAIT cells, where a small subset showed clear upregulation of genes involved in TCR signaling (*NR4A1*, *NFKBID*, *REL*; MAIT_c3, Supplementary Fig 3D). Fifth, unlike T_conv_, in the thymus we discovered a proportion of both iNKT and MAIT cells having an effector signature. We found a cluster of iNKT cells (NKT_c6) expressing classically iNKT-associated genes along with upregulation of effector genes usually associated with type I immunity, such as *EOMES* and *GZMK* (Fig 2M, Supplementary Fig 3C). Some of these cells expressed CD8A transcripts, suggesting that CD4^+^ and CD8^+^ iNKT cells might develop into transcriptionally distinct subsets, with CD8^+^ iNKT cells having a more effector-associated signature. For MAIT cells, we found a similar pattern, with cells expressing genes previously associated with MAIT cells (*KLRB1*, *SLC4A10*, *IL23R*) (Dusseaux et al., 2011; Park et al., 2019) also displaying an effector transcriptome signature (MAIT_c6, Supplementary Fig 3D). This was evidenced by the upregulation of genes encoding for granzymes (*GZMA* and *GZMK*), chemokines (*CCL5*), chemokine receptors (*CCR6*), and transcription factors usually associated with type I (*EOMES*) or type 3 (*RORA*) immunity (Fig 2M and Supplementary 8). In our integrative analysis from both the thymus and blood, we identified a shared utilization of these effector programs by iNKT, MAIT and γδ T cells. Specifically, approximately 28.7% ± 22.3% of thymic T_inn_ cells displayed an effector signature, as depicted in Fig 1G, I (clusters 12-17). To delve deeper into whether these effector T_inn_ cells in the thymus are cells that initially acquired an effector signature in the blood and subsequently recirculated back to the thymus, we conducted a comparative examination of gene expression profiles between effector iNKT and MAIT cells in the thymus and blood (Supplementary Fig 5C). Our results revealed distinctive tissue-specific gene expression profiles for both cell types. Thymic cells exhibited higher expression levels of genes such as *CCR7*, *TOX*, *TOX2*, and *SOX4*. Conversely, blood-derived cells demonstrated elevated expression of genes including *DUSP2* and *BCL2*. These findings strongly suggest that thymic T_inn_ cells found in the effector clusters possess a unique transcriptome when compared to their blood counterparts. Therefore, it is unlikely that these effector T_inn_ cells are derived from recirculating cells originating in the blood.

**Figure 2:**
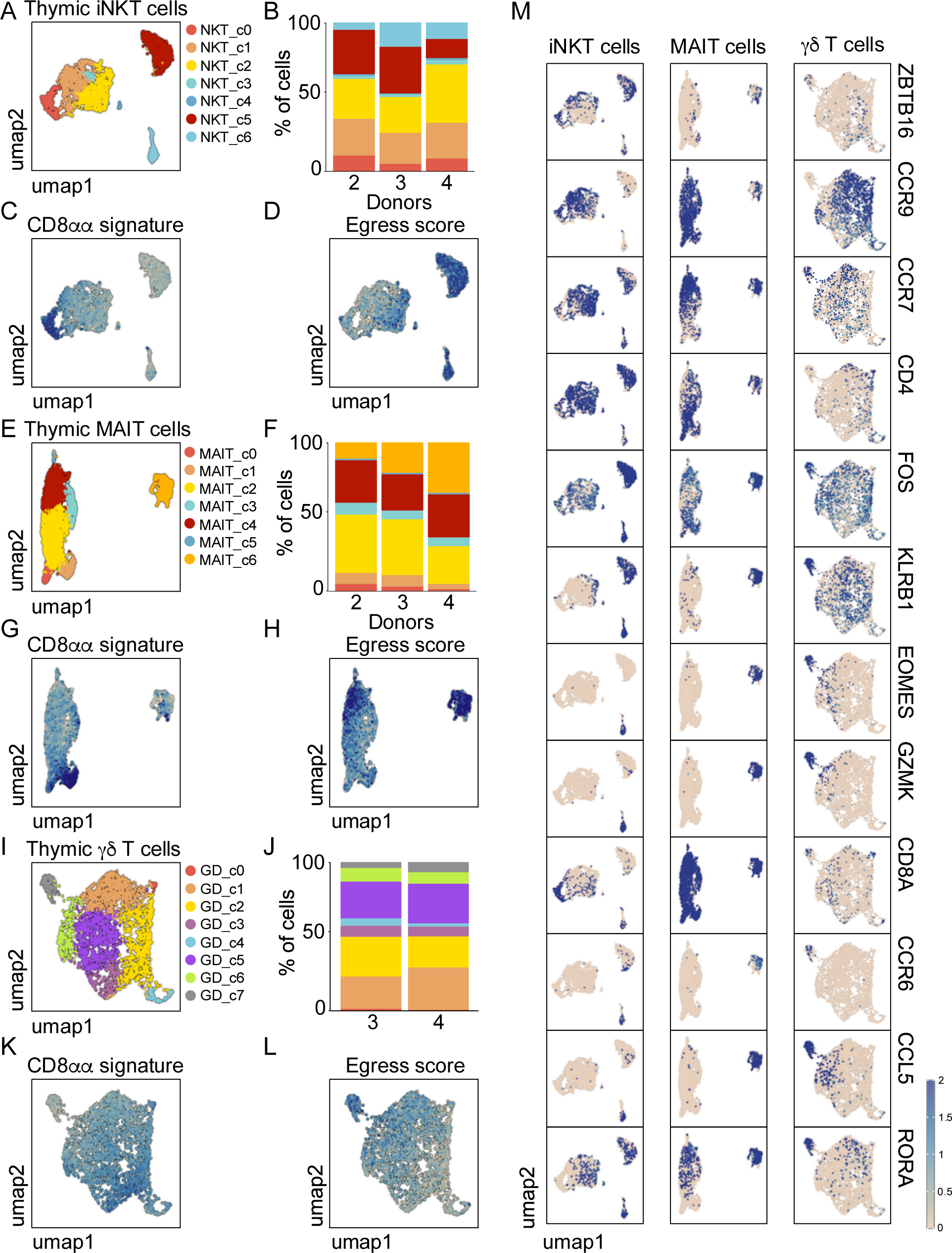
Human Innate T cell development. Clustering of hashtag-separated thymic iNKT (A), MAIT (E) and γδ T cells (I) and the respective proportion of cells per cluster and donor (B, F, J), the projection of the CD8αα signature (C, G, K) and egress score (D, H, L). M. Characteristic gene expression in thymic iNKT, MAIT and γδ cells. n=4 postnatal human thymus samples for all panels.

Finally, we also re-analyzed post-natal thymic human γδ T cells separately and identified 8 transcriptionally distinct clusters (Fig 2I and Supplementary Fig 3E, Table VII), with findings largely replicating a recent report on pediatric γδ thymocytes (Sanchez Sanchez *et al*., 2022). We found immature populations observed in GD_c0, 1, and 2, cells with TCR activation/co-stimulation profiles (GD_c3), type I interferon response signature (GD_c6) and effector γδ T cells displaying an Egress gene signature and a mixed type 1/type 3 effector potential (GD_c7). We also observed cells with a cycling gene signature (GD_c4), which was notably absent in iNKT and MAIT cells. Overall, our findings demonstrate that only a small proportion of T_inn_ cells in the thymus exhibit a transcriptional signature associated with an effector program, and that this effector program has a distinct mixed type 1/type 3 effector potential.

### Effect of clonal selection on iNKT, MAIT and γδ T cells effector states

To investigate whether the T_inn_ cells exhibiting an effector transcriptome possessed a distinct TCR repertoire compared to the naïve T_inn_ cells identified in the human thymus, we conducted paired VDJ sequencing (Fig 3). This allowed us to link the different cell states to their corresponding TCR sequences, providing insights into the diversity and specificity of the TCR repertoire within these distinct T_inn_ cell populations. As a measure for TCR diversity, we used the Shannon index comparing naïve-like cells to effector-like cells, as determined by the cluster assignment on the re-analyzed cell types described above. For iNKT cells, we found that most of the VDJ sequenced cells used the *TRAV10* gene segment rearranged with *TRAJ18* (Fig 3A, B), resulting in a CDR3 size of 14 amino acids with a canonical sequence (Fig 3C), emphasizing the importance of the CDR3α for antigen recognition by iNKT cells (Scott-Browne et al., 2007). This invariant TCRα chain was paired with diverse TCRβ rearrangements involving primarily the *TBV25* chain (Fig 3D), which was evenly used across all clusters (Fig 3E). We did not observe any shared TCR clonotypes between the naive- and effector-like cells. Additionally, the Shannon indexes for TCR clonotypes were found to be identical across both naïve- and effector-like cells, suggesting the absence of clonal selection associated with the development of an effector transcriptome.

**Figure 3:**
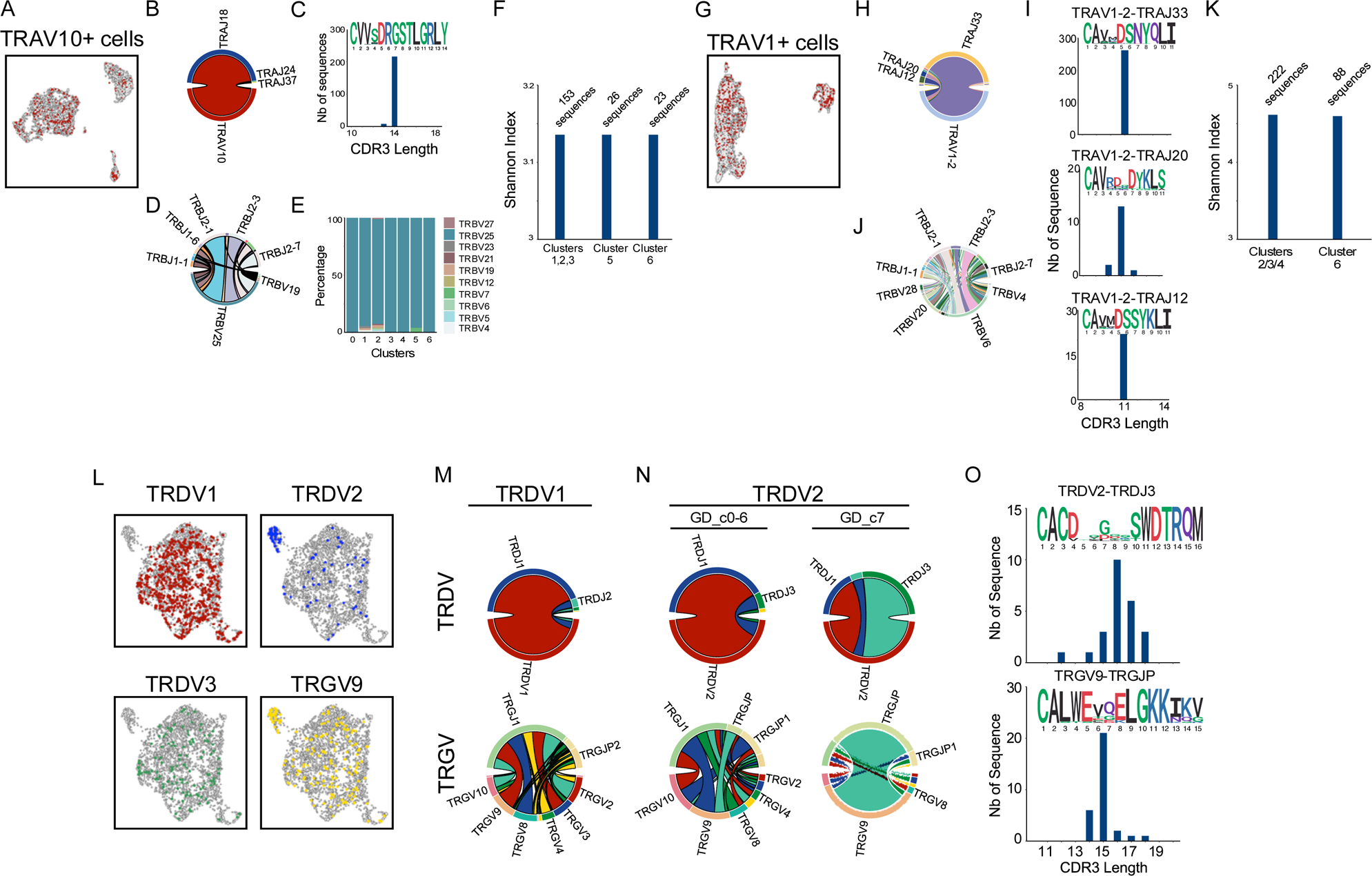
Innate T cell TCR diversity during development. Cells with VDJ sequencing and their cell-type specific characteristic chain arrangement for thymic iNKT (A), MAIT (G) and γδ T cells (L). For each cell type, the respective proportions of gene segment usage in each chain (B, D; H, J; M, N) are shown together with their CDR3 length and sequence logo (C, I, O) and their cluster-specific usage (E, with clusters as in Fig 2A). Shannon Index as an estimation of TCR diversity in the naive-like and effector-like iNKT (F) and MAIT (K) cells, based on clusters in Figure 2A and E, respectively. n= 1 human thymus sample for panels A-O.

Thymic MAIT cells largely used the *TRAV1-2* gene segment rearranged primarily with *TRAJ33*, *TRAJ20* and *TRAJ12* (Fig 3G, H), as previously reported (Reantragoon *et al*., 2013). These rearrangements were largely of the same CDR3 size with limited sequence diversity and notably, contained the conserved Y95 residue within the CDR3α loop (Fig 3I), which is crucial to MAIT cell activation (Reantragoon et al., 2012; Young et al., 2013). These TCRα chains were paired with a diverse repertoire of TCRβ chains (Fig 3J), dominated by the usage of the *TRBV6*, *TRBV20* and *TRBV4* gene segments (Reantragoon *et al*., 2013; Tilloy et al., 1999). Similar to our observations with iNKT cells, we found no sharing of TCR clonotypes and no evidence of clonal selection among MAIT cells with an effector transcriptome (MAIT_c6) compared to naïve-like cells (MAIT_c2-4) based on the Shannon index of TCR clonotypes (Fig 3K). In contrast, effector γδ T cells (GD_c7) were enriched for cells expressing the *TRDV2* and *TRGV9* gene segments, while cells expressing *TRDV1* and *TRDV3* gene segments were excluded from this cluster (Fig 3L). However, some *TRDV2*^+^ or *TRGV9*^+^ cells could also be found in the non-effector clusters, suggesting a potential role for these gene segments in the development of effector γδ T cells in the post-natal human thymus. Supporting this hypothesis, we observed that the rearrangements of both the Vδ2 chains and associated Vγ9 chains differed largely between cells in the effector versus non-effector clusters (Fig 3N). Specifically, the Vγ9 chains of effector cells were found to be preferentially rearranged with the TRGJP gene segment and enriched for the public CDR3 sequence typically found amongst Vδ2Vγ9 γδ T cells in the adult blood (Davey et al., 2018) (Fig 3O), whereas Vδ2^+^ cells in the non-effector clusters showed more diverse Vγ gene usage and rearrangements (Fig 3N). In summary, the acquisition of the effector programs in iNKT and MAIT cells is not associated with changes in TCR diversity, while the rearrangements of Vδ2 and Vγ9 chains in γδ T cells suggest predisposition towards the effector program.

### Gene Expression Programs that Characterize T cell Effector Functions

Our detailed analysis of thymic T cell populations and gene expression modules revealed shared developmental patterns between iNKT, MAIT, and T_conv_ cells. To further characterize the functionality of T_inn_ and T_conv_ cells in the blood, we initially examined the distribution of these cell types across transcriptional clusters. In the blood, conventional CD4^+^ T cells were primarily located in cluster 11 (representing naïve CD4 T cells) and 7 (comprising Tregs). However, a variable proportion of CD4^+^ T cells was also observed in effector clusters, particularly clusters 12 and 16, and this proportion varied among donors, ranging from 18.4% to 93.7% of cells (Fig 1G & J). Blood CD8^+^ T cells were predominantly found in cluster 10, representing naïve CD8 T cells, although varying proportions of CD8^+^ T cells were present in effector clusters. This variability may reflect differences in the immunological history of each donor, with proportions ranging from 16.5% to 94% of cells (Fig 1J). In striking contrast, the majority of blood T_inn_ cells (approximately 94.3% ± 6.7%) were distributed across effector clusters 12 to 17, irrespective of the donor (Fig 1J).

We next investigated the transcriptional states in blood T cell populations using the cell hashtags to reanalyze blood iNKT, MAIT, γδ T cells and T_conv_ CD4^+^ and CD8^+^ T cells individually (Fig 4A, B). Our analysis, utilizing the previously identified naïve and effector gene modules (GEP3-6; Fig 1K, Supplementary Tables VIII-XII), indicated that each of the investigated cell types could be found within these identified gene programs, albeit with varying proportions for each cell type (Fig 4C, D). To provide further context and understanding of these gene modules, we computed overlap scores and statistically assessed their enrichment with literature-derived signatures (Cano-Gamez et al., 2020; Poon et al., 2023; Rose et al., 2023; Terekhova et al., 2023). Subsequently, we scored the joint signature-GEP interactions in our dataset (Supplementary Fig 9). GEP3 was found to be closely associated with signatures of naïve T cell characteristics. In contrast, GEP4 displayed similarities with central memory T cells (T_cm_), effector memory (T_em_) or literature-derived signatures classified as a mix thereof (T_cm_/T_em_), while GEP6 exhibited characteristics akin to terminally differentiated effector memory cells (T_emra_). GEP5, on the other hand, shared elements with T_em_ cells and previously identified CD8 MAIT signatures (Supplementary Fig 9A). When examining blood iNKT cells, we noticed that they predominantly fell into two categories, expressing either the GEP3 or GEP5 programs (Fig 4D). However, there was also a subset of iNKT cells that exhibited either the GEP4 or GEP6 programs. Interestingly, the distribution of these programs varied significantly among different donors (Fig 4B). Notably, iNKT cells characterized by the GEP4 program expressed CD4 transcripts, whereas those using GEP5 or GEP6 had lost CD4 expression (Supplementary Fig 10A). To validate this observation, we examined the cellular phenotype of blood iNKT cells. We found that blood CD4^+^ iNKT cells were mostly PLZF^−^ CD161^−^ EOMES^−^ GZMK^−^ but CCR7^+^, suggesting that they likely belonged to the naïve GEP3 program (Supplementary Fig 10). In contrast, CD8^+^ and DN iNKT cells were mostly PLZF^+^ CD161^+^ and displayed an effector phenotype (EOMES^+^ GZMK^+^ CCR7^−^ CD62L^−^, Supplementary Fig 10). These findings are in line with previous data indicating that CD4-negative iNKT cells become more prevalent in the blood with age, eventually becoming the dominant population in the adult blood iNKT cell compartment (Berzins et al., 2005; Sandberg et al., 2004). This suggests that CD4-negative iNKT cells may originate from CD4^+^ iNKT cells that undergo a loss of CD4 expression (and potentially gain CD8 expression in some cases) as they transition toward a more effector-like state. Conversely, when examining MAIT cells in the blood, we observed that the majority of them exhibited the GEP5 program, with only a minor fraction utilizing the GEP6 program (Figure 4D). This GEP phenotype was further confirmed by flow cytometry, revealing that most MAIT cells were CD8^+^, and interestingly, all MAIT cells displayed a uniform effector state (characterized by PLZF^+^ CD161^+^ EOMES^+^ GZMK^+^ CCR7^−^ CD62L^−^), regardless of CD8 expression (Supplementary Figure 11). These findings indicate that MAIT cells in the bloodstream primarily exist in a single cell state, aligning with recent data demonstrating minimal transcriptional heterogeneity among blood and liver-resident MAIT cells (Garner *et al*., 2023), and no significant transcriptional distinctions between CD8^+^ and DN MAIT cells.

**Figure 4:**
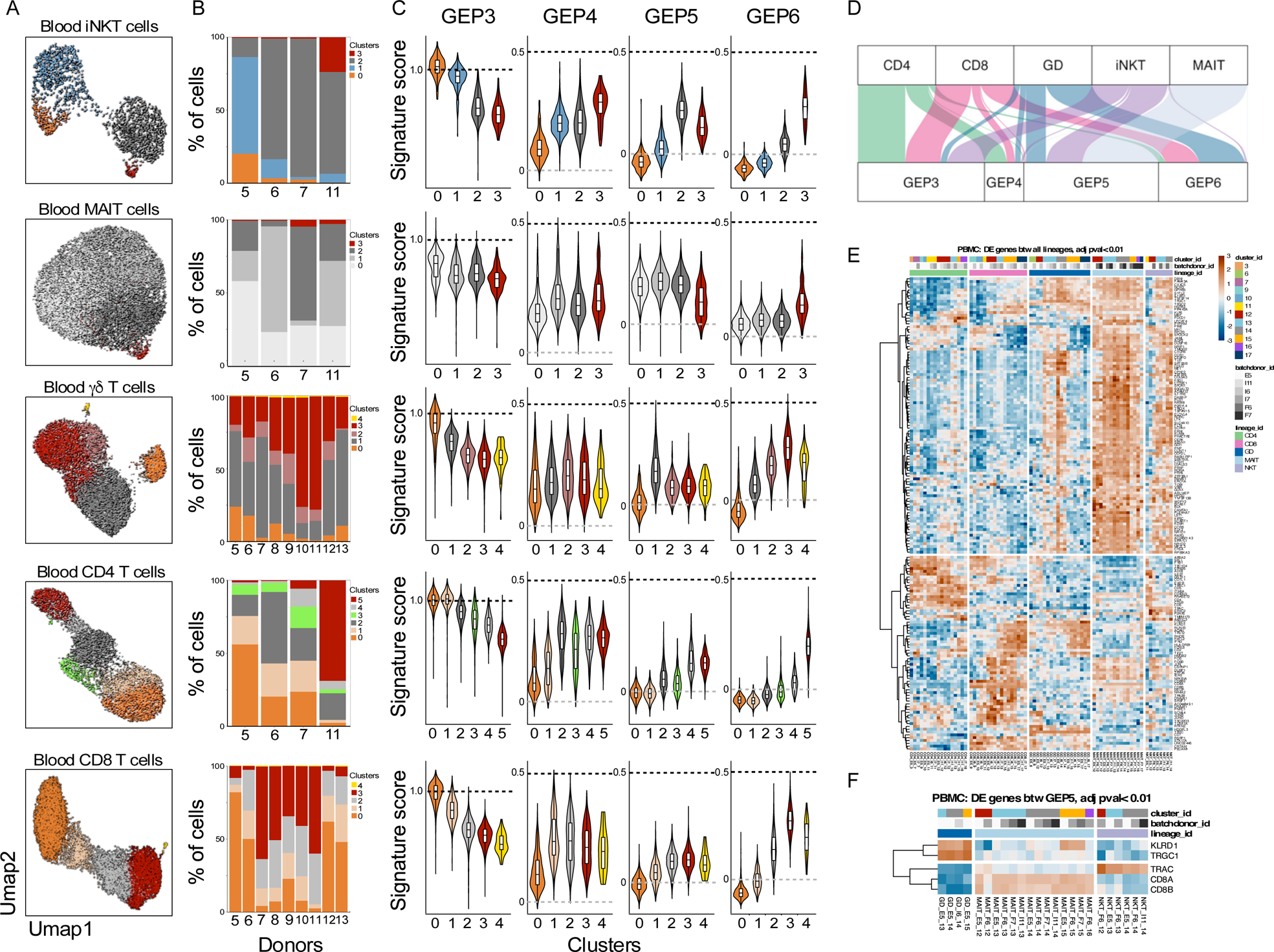
Gene expression programs in T_inn_ and T_conv_. A. Clustering of hashtag-separated blood iNKT, MAIT, γδ T cells, CD4 and CD8 T cells, B. the respective proportion of cells per cluster and donor and C. the effector Gene expression program (GEP) signature scores (as in Fig 1K) per cell type and cluster. D. GEP usage for each cell type, based on cNMF-derived usage matrix. E. Pseudo-bulk, pair-wise differential gene expression between cell-types (upper panel). T_inn_ specific signature genes in GEP5 (lower panel). For both panels, genes with p < 0.01 are depicted. n=4,4,9,4,9 for iNKT, MAIT, γδ, CD4 and CD8 cells respectively.

Blood γδ T cells were stratified into five distinct clusters, with one cluster corresponding to naïve cells (c0, GEP3) and another cluster (c4, GEP11) representing cycling cells. The majority of cells in clusters c1-c3 were categorized into either the GEP5 or GEP6 program (Figure 4C, D), and this division in GEP utilization closely mirrored the specific TCR usage among these cells. Specifically, TRDV2/TRGV9-expressing cells were predominantly associated with the GEP5 program, whereas cells expressing TRDV1^+^ or TRDV3^+^ were enriched in clusters expressing GEP6 (Supplementary Fig 12A-C). These GEP phenotypes were further validated through flow cytometry, revealing that Vγ9^+^Vδ2^+^ T cells primarily expressed PLZF and GZMK, while Vδ2^−^ T cells were PLZF^−^ but instead GZMB^+^ (Supplementary Figure 12D, E). In light of this, it appears that the GEP5 program represents an effector gene module exclusively expressed by innate T cells, suggesting that human T_inn_ cells share a common transcriptional state. This observation parallels the way mouse iNKT and MAIT cells share type 1 or type 3 immunity effector states (Krovi *et al*., 2022). Regarding the distribution of CD4 and CD8 T_conv_ cells in the blood, it revealed two primary patterns. Some T_conv_ cells were found within clusters containing naïve cells (clusters 0 and 1), characterized by high expression of GEP3. In contrast, others were dispersed across clusters of cells displaying a gradient of the GEP6 program, with intermediary cells expressing GEP4 (Supplementary Figure 13). The proportions of cells in these clusters exhibited variations among donors (Fig 4B). In summary, these findings underscore the distinct associations between different T cell types and effector programs. In the blood, iNKT, MAIT, and Vγ9^+^Vδ2^+^ γδ T cells predominantly employ the GEP5 program, a program also shared by effector T_inn_ cells in the thymus (Supplementary Fig 7). Conversely, conventional CD4^+^ and CD8^+^ T cells transition into effector cells along a gradient defined by the GEP6 program. Notably, this GEP6 program is also shared by Vδ3^+^ and Vδ1^+^ γδ T cells.

Next, we aimed to identify genes specific to each T cell lineage in human blood. We conducted pairwise DEG analyses between the lineages using a pseudo-bulk method in conjunction with the Likelihood Ratio Test after accounting for batch effects. As a result, we uncovered a total of 167 genes that exhibited significant differential expression (padj < 0.01) in at least two of the comparisons we conducted (Figure 4E). These distinct patterns of differential expression provided insights into changes linked to the transition from a “naive” state to an “effector” state across various cell types. Furthermore, we identified genes that were commonly expressed by two distinct cell types when compared to the others. Nevertheless, we did not readily discern any gene expression patterns specific to a particular cell type. However, intriguingly, among these, a group of 104 genes appeared to be capable of distinguishing γδ, MAIT, and iNKT cells from T_conv_ CD4 and CD8 T cells. It is noteworthy that 63% of these genes overlap with the previously identified GEP5 program. Given that only Vγ9^+^/Vδ2^+^ T cells share the GEP5 program with iNKT and MAIT cells, while Vδ2^−^ T cells exhibit greater similarity to T_conv_ cells as they share the GEP6 program (Supplementary Fig 12), we explored whether we could identify cell-type-specific gene signatures specifically among GEP5-expressing cells, using the same analytical approach. Surprisingly, the results demonstrated that the only genes expressed at significantly different levels across iNKT, MAIT, and γδ T cells when employing the GEP5 program (Fig 4F) were genes encoding the constant regions of the TCR genes (*TRGC1*, *TRAC*), the CD8 coreceptor (*CD8A*, *CD8B*), and the CD94 receptor (encoded by *KLRD1*). These findings collectively suggest that in the human blood, T_inn_ cells, which encompass iNKT, MAIT, and Vγ9^+^/Vδ2^+^ cells, distinguish themselves from T_conv_ cells by employing a specific gene program, but there is minimal transcriptional difference among T_inn_ cells themselves.

### The effector GEPs exhibit distinct migration, cytokine, chemokine and integrin characteristics established by distinct Gene Regulatory Networks

The differentiation states of T cells are intricately tied to their phenotypic, functional, and migratory attributes, rendering the characterization of these states highly relevant from a clinical perspective. In fact, our findings underscore that each GEP is aligned with the preferential expression of distinct sets of chemokine and cytokine receptors, as well as chemokines, cytokines, cytotoxicity-related molecules, NK receptors, and integrins (Fig 5A). To exemplify, the GEP4 program, shared notably by both T_cm_/T_em_ (Supplementary Fig 9) and, to some extent, iNKT cells depending on the donor (Fig 4B-D), demonstrates heightened transcription levels of genes coding for the chemokine receptors CXCR3 and CCR4, the Sphingosine-1-Phosphate Receptor 4, and the oxysterol receptor GPR183. The latter has been documented to offer survival and migratory signals to thymocytes and CD4 T follicular helper cells(Li et al., 2016). Furthermore, the GEP4 program showcases heightened expression levels of *IL2RA*, *IL6R*, *IL-4R*, and the integrin *ITGB1*, while conspicuously lacking transcripts linked to cytotoxic molecules. Conversely, the GEP5 program, closely linked to the majority of cells from T_inn_ cell subsets (iNKT, MAIT, Vδ2γ9^+^ γδ T cells) across most donors (Fig 4B-D), demonstrates elevated expression of diverse chemokine receptors, including *CCR1*, *CCR2*, *CCR5*, *CCR6*, as well as *CXCR6* and of cytokine receptors such as *IL18R1*, *IL18RAP*, *IL12RB1*, *IL12RB2*, *IL23R*, and *IFNGR1* (Fig 5A). This distinct gene expression pattern is further marked by the presence of granzymes A and K transcripts, while granzymes B and H are notably absent (Fig 5A and Supplementary Fig 5A). Another noteworthy hallmark of the GEP5 program is the expression of the NK receptor *KLRB1*. These findings closely align with previously identified markers associated with human MAIT and Vδ2γ9^+^ γδ T cells (Davey *et al*., 2018; Meermeier et al., 2022; Park *et al*., 2019) and are consistent with the demonstrated ability of T_inn_ cells to respond to inflammatory cytokines like IL-12, IL-18, and IL-23, even without TCR engagement. On the other hand, the GEP6 program, primarily associated with T_em_/_emra_ cells (Supplementary Fig 9) and Vδ1^+^ and Vδ3^+^ γδ T cells across the majority of donors (Fig 4B, C and Supplementary Figure 12), is characterized by increased expression of CX3CR1 (Supplementary Figure 13). The graded expression of CX3CR1 correlates with the differentiation of both CD4 and CD8 T cells towards an effector state (Zwijnenburg et al., 2023). Additionally, the GEP6 program exhibits heightened expression of transcripts encoding *IFNG*, *CCL4*, *CCL5*, *KLRD1*, as well as several integrins (*ITGAL*, *ITGB2*, *ITGAM*). It also includes genes associated with cytotoxicity, such as granzymes B, H, and granulysin (*GNLY*). Interestingly, transcripts for granzyme K are significantly reduced in GEP6 compared to GEP5. These results are in line with recent findings indicating that GZMK^+^ and GZMB^+^ cells delineate T_cm_ and T_em_/T_emra_ T cell populations (Duquette et al., 2023; Jonsson et al., 2022).

**Figure 5:**
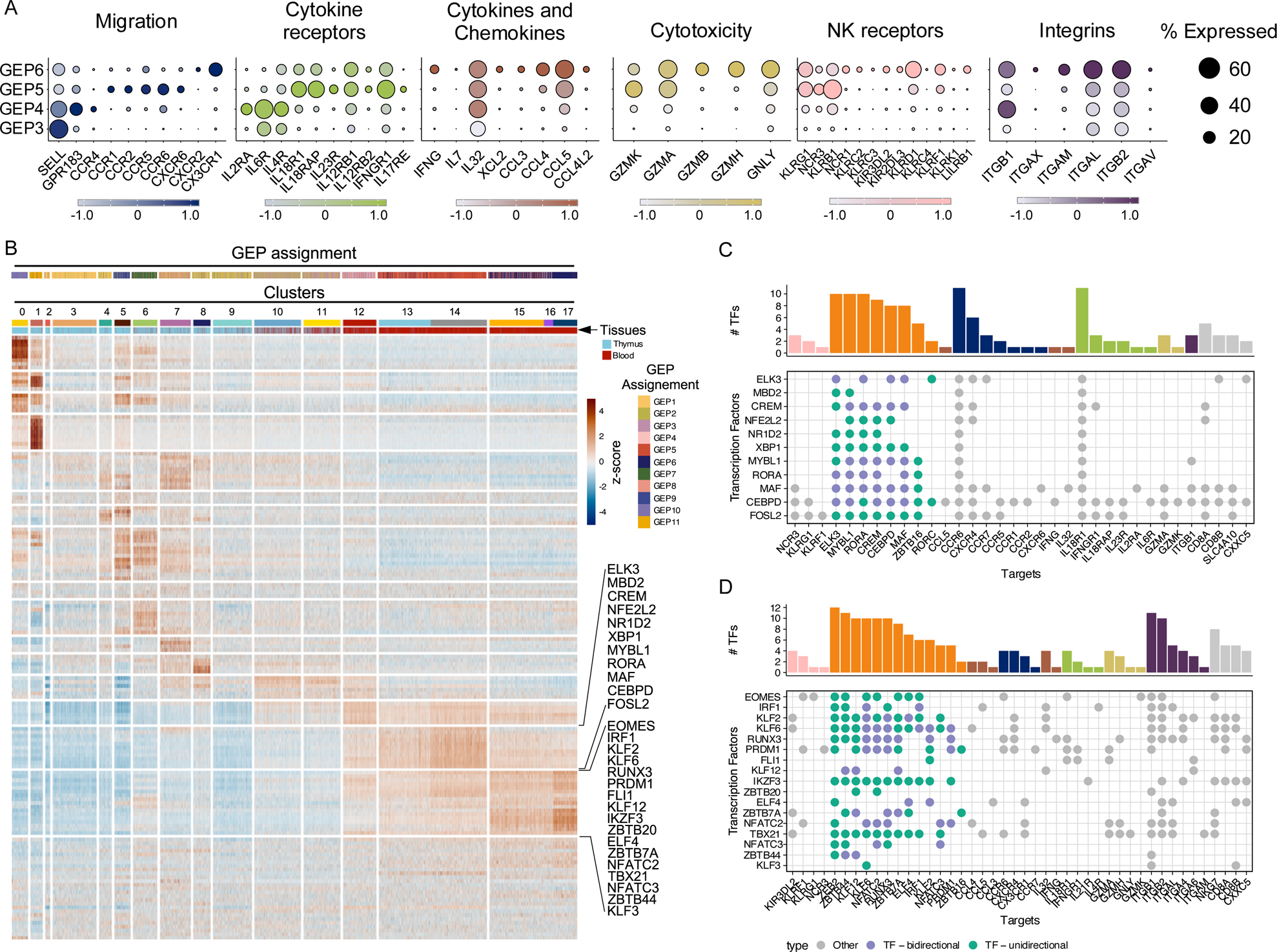
Effector gene expression programs in T_inn_ and T_conv_. A. Key genes categorized by function and depicted by their level (z-score color scale) and percentage of expression in cells belonging to the indicated GEPs. B. Single-Cell Regulatory Network Inference and Clustering of transcription factors (TFs) and their expression strength per cell (as row-scaled z-scores), ordered by cluster (as in Fig 1C), with tissue of origin and GEP assignement based on cNMF usage indicated by color bar. Two row clusters are marked which are preferentially enriched in T_inn_ (upper bracket) and T_conv_ (lower bracket) C./D. TFs with pronounced activity in (C) T_inn_ and (D) T_conv_ (corresponding to brackets in B) and their targets. Green indicates TF (y-axis) that have other TFs as their target (x-axis), where purple labels TFs that can interact in either direction. The marginal bar chart shows the number of TFs per target, color-coded by their functional categorization (as in A).

To predict gene regulatory networks associated with these gene programs, we used Single-Cell Regulatory Network Inference and Clustering to identify enriched TFs with their direct downstream targets and scored the activity of these so-called regulons in single cells (Aibar et al., 2017). We identified a total of 149 regulons that were expressed in at least 20% of the cells within a specific cluster and displayed associations with different clusters (Fig 5B). Notably, 11 regulons exhibited more pronounced activity in T_inn_ cells compared to T_conv_ cells (Fig 5B). These regulons were governed by TFs such as *ELK3*, *MBD2*, *CREM*, *NFE2L2*, *NR1D2*, *XBP1*, *MYBL1*, *RORA*, *MAF*, *CEBPD*, and *FOSL2*. Curated analysis of their predicted target genes indicates that these transcription factors may play a central role in shaping the unique transcriptional profile observed in T_inn_ cells during steady-state conditions. This role potentially encompasses the regulation of chemokine and cytokine receptors, as well as other genes associated with T_inn_ cells, including ZBTB16 (which encodes PLZF), the master regulator of the T_inn_ cell lineage (Fig 5C). Interestingly, previous data established CEBPD as a regulator of CCR6 expression in human MAIT cells(Lee et al., 2018). Furthermore, *FOSL2* (encoding Fra2) has been implicated in the normal development of mouse iNKT cells(Lawson et al., 2009), and c-Maf has been recognized as a key player in the differentiation of IL-17-producing mouse iNKT cells (Yu et al., 2017). Additionally, RORA has been described as an auxiliary transcription factor for Th17 cells (Ciofani et al., 2012), and there is evidence of CREM, XBP-1 and NR1D family of TFs involvement in the regulation of Th17 cells as well (Chang et al., 2019; Yoshida et al., 2016). The remaining transcription factors in this list, to the best of our knowledge, have not been extensively studied in the context of T_inn_ cell development and/or function. In addition to this set of regulons, another group of regulons exhibited enriched activity within effector T_conv_ cells, although some shared activity with T_inn_ cells (including *EOMES*, *RUNX3*, *PRDM1*, and *FLI1*; Fig 5B). These findings are consistent with previous results showing that EOMES and RUNX3 collaborate to promote the formation of the transcriptional program through epigenetic programming of innate memory CD8 T cells in mice (Istaces et al., 2019). This suggests that similar mechanisms might be at play in human T_inn_ cells. As T_conv_ cells differentiate into T_em/emra_ cells, there is an increased activity of regulons driven by *TBET*, *KLF*, and *NFAT* family transcription factors (Fig 5B and D), in agreement with their functions in regulating the cytolytic activity of CD8 T cells (Intlekofer et al., 2005; Klein-Hessling et al., 2017; Nah and Seong, 2022). Taken together, our data unveil novel candidate regulators of T_inn_ and T_conv_ effector programs, along with their predicted target genes, which warrant further experimental validation.

### Cross-species analysis of thymic T_inn_ cells development

Our analysis of human T_inn_ and T_conv_ cells across thymus and bloodstream revealed common transcriptional patterns during their development. Interestingly, in the thymus, only a minority of T_inn_ cells displayed an effector phenotype, marked by a unique gene expression program that we named GEP5. This differs from the predominant effector association of T_inn_ cells in mouse thymus, where distinct effector subsets that closely resemble CD4 T helper cells and innate lymphoid cells develop and reside (Krovi *et al*., 2022; Lee et al., 2020; Legoux et al., 2019). To explore the similarities in the transcriptional signatures of mouse and human T_inn_ cells, we first constructed a reference mouse T_inn_ dataset, comprising data from nine different studies (Baranek et al., 2020; Chandra *et al*., 2023; Harsha Krovi et al., 2020; Koay et al., 2019; Lee *et al*., 2020; Legoux *et al*., 2019; Li et al., 2022; Maas-Bauer et al., 2021; Wang et al., 2023). This merged dataset revealed 13 transcriptionally distinct clusters, where iNKT, MAIT, and γδ T cells coexisted, with variable proportions (Fig 6A and Supplementary Fig 14, Table XIII). In addition, there are also lineage-specific clusters, like clusters 1 and 2 unique to γδ T cells, representing immature *Cd24a*^+^ *Gzma*^+^ cells (Lee *et al*., 2020; Li *et al*., 2022). Other clusters represented signaling cells committing to innate T cell lineage (in clusters 3 and 4), cycling cells (in cluster 5), type I cells in clusters 8 and 9, type II cells in clusters 6 and 7 and type III cells in clusters 10 and 11. Cluster 0 expressing *Sell* (encoding for Cd62l), *Klf2*, *Ccr7*, *Foxo1* and *S1pr1* and associated with post-selection/naive T cells likely represents T_inn_ cells positively selected on thymic epithelial cells (TECs), bypassing the “innate” pathway (Krovi *et al*., 2022; Salou et al., 2021). Altogether, this analysis enabled us to establish the fundamental signatures of mouse T_inn_ subsets and to perform a cross-species comparison of cell identities. Using a meta-analysis approach that compares similarities between cell clusters, we assessed the pairwise correspondence between these murine T_inn_ signatures and our human iNKT, MAIT, and γδ T cell clusters (Fig 6B). Among the human iNKT cells residing in cluster 0 (NKT_c0) and exhibiting a CD8αα T cell gene signature, we observed the strongest resemblance to the signaling cells (Fig 6B). This similarity is likely due to the shared expression of numerous genes associated with TCR activation. Conversely, cells with an effector profile (NKT_c5 and NKT_c6) showed the closest relationship to mouse type I and type III cells (Fig 6B). Importantly, we did not find human clusters corresponding uniquely to specific mouse subsets, confirming that human iNKT cells do not differentiate into distinct subsets but rather acquire a mixed type I/type III transcriptome. We also did not detect any human iNKT cell clusters that matched with the mouse type II subset with a high degree of confidence (AUROC > 0.65; Fig 6B), suggesting that type II iNKT cells are likely absent in the human thymus. Corroborating this finding, we did not detect any expression of IL-4 or IL-13-encoding transcripts in human thymic iNKT cells, which are typically associated with mouse type II thymic iNKT cells. Similar patterns were observed for MAIT and γδ T cells in the human thymus, with effector cells resembling mouse type I and type III effector cells (Fig 6B) indicating that a limited subset of T_inn_ cells in humans follows a distinctive path, displaying mixed effector potential, unlike the mouse model where multiple effector subsets are identified. We next assessed whether the TFs responsible for driving the characteristic regulons of human T_inn_ cells (Fig 5A) are also expressed in mouse T_inn_ cells (Fig 6C). Our analysis revealed that most of these TFs are indeed expressed in mouse T_inn_ cells as well, although their distribution of expression across clusters varies. These results suggest that each TF may have a unique role in shaping the distinct subsets of mouse T_inn_ cells. They also raise the possibility of some similarities in transcriptional regulation between the two species during the development of T_inn_ cells’ effector functions in the thymus. However, there are exceptions. Notably, while *CEBPD*, *EOMES*, and *MYBL1* are highly expressed in human T_inn_ cells (Supplementary Fig 5A), their expression in mouse T_inn_ cells is barely detectable (Fig 6C). On the other hand, mouse type I T_inn_ cells exhibit high levels of Tbet (encoded by *Tbx21*, Fig 6C), whereas human T_inn_ cells have low Tbet expression (Supplementary Fig 5A). These findings highlight some species-specific differences in TF expression that could play a role in modulating T_inn_ cell development and functions.

**Figure 6:**
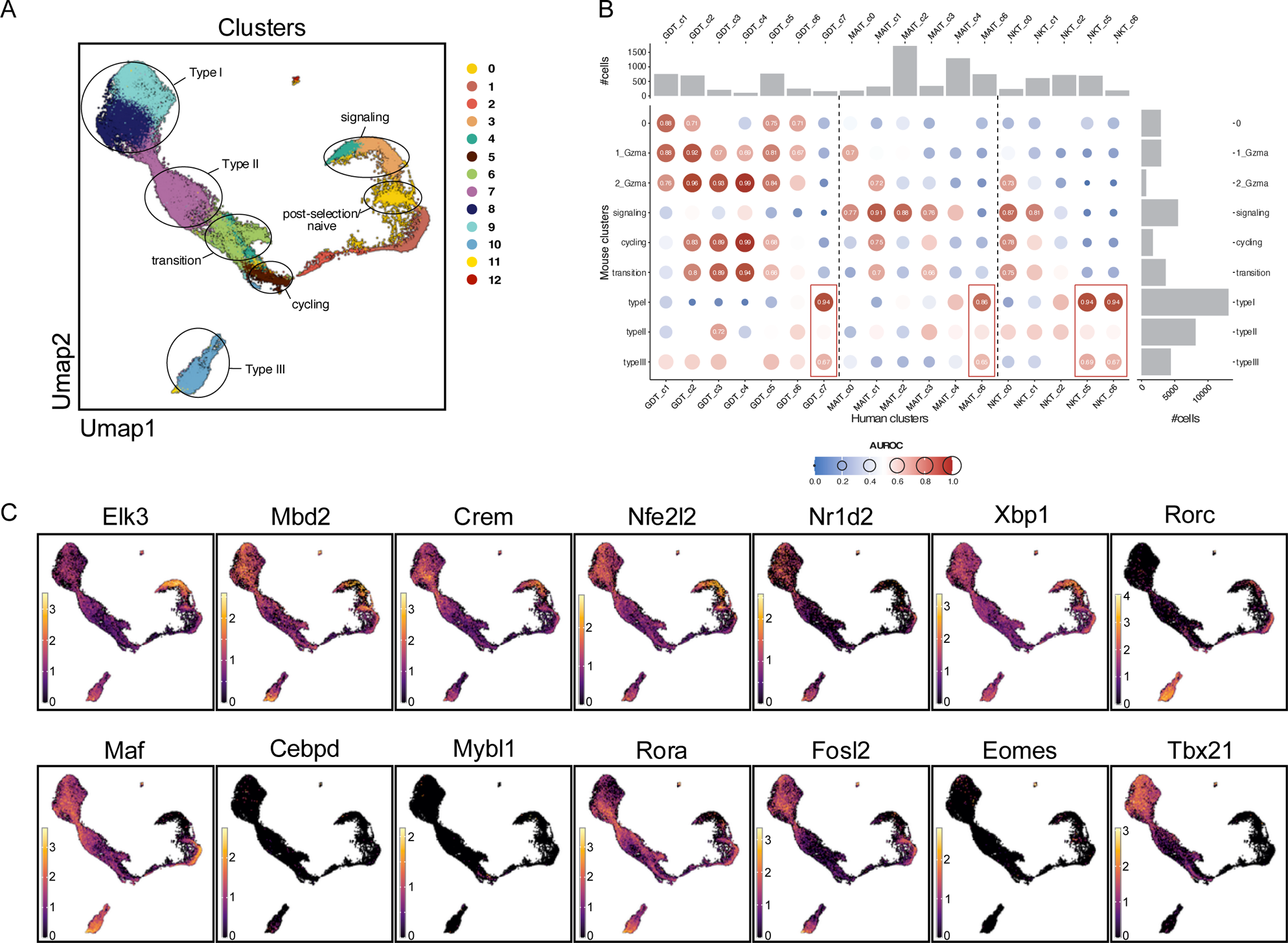
Cross-species comparison of mouse and human T_inn_ development. A. Mouse T_inn_ reference atlas with 7 characteristic cell states highlighted that are found across lineages (as in Supplementary Figure 14). B. Meta-neighbour analyses showing pairwise correspondence (AUROC scores) between murine T_inn_ (as in A) and human iNKT, MAIT and γδ T cell clusters (as in Figure 2). Marginal bar charts indicate number of cells in the corresponding clusters. C. Expression of human regulon-driving transcription factors (as in Figure 5) together with murine TFs of importance in T_inn_ development (Rorc, Tbx21) projected on mouse T_inn_ reference atlas (as in A.)

### CD1D expression in the mouse and human thymus

The existence of T_inn_ cells displaying a transcriptome akin to developing T_conv_ cells in the human thymus raises questions about their origin. In mice, a subset of MAIT cells can be positively selected by radiation-resistant TECs (Chandra *et al*., 2023; Legoux *et al*., 2019). In such instances, MAIT cells do not follow the usual path of acquiring a memory or effector phenotype, which happens when they are positively selected by DP thymocytes (Krovi *et al*., 2022). This is because TECs lack the expression of SLAM receptors, which serve as crucial secondary signals for T_inn_ commitment (Griewank et al., 2007). Although such naïve cells are more common among MAIT cells, a small subset of thymic mouse iNKT cells exhibit a similar transcriptome (Krovi *et al*., 2022). We hypothesized that the presence of naïve T_inn_ cells in humans might be attributed to a similar process involving TEC-mediated selection. Given that MR1-encoding transcripts are broadly expressed and surface MR1 expression can be challenging to detect, we opted to investigate the expression of CD1D protein in the thymus instead. Mouse TECs were previously reported to express CD1d on their surface (Forestier et al., 2003). Analysis of transcripts encoding Cd1d1 from the mouse thymus single-cell RNA sequencing atlas (Park *et al*., 2020) confirmed expression across various cell types, including all thymocyte populations as well as cortical and medullary thymic epithelial cells (Fig 7A, B). These findings were further corroborated through flow cytometry analyses (Fig 7C, D). In contrast, analyses using data from the human thymus single-cell RNA sequencing atlas(Park *et al*., 2020) revealed a more limited pattern of *CD1D* expression (Fig 7E, F). Consistent with the mouse data, human DP thymocytes express transcripts encoding CD1D and have CD1D molecules on their surface, but this expression is lost in mature single-positive thymocytes. Additionally, while human cortical thymic epithelial cells (cTECs) express transcripts encoding *CD1D* and have CD1D protein on their cell surface, medullary thymic epithelial cells (mTECs) do not exhibit this expression (Fig 7G, H). Interestingly, the crucial role of mTECs in the intra-thymic development of murine iNKT cells has been established (Cui *et al*., 2022; White et al., 2014), suggesting that this interspecies difference in CD1D expression may affect iNKT cell development.

**Figure 7:**
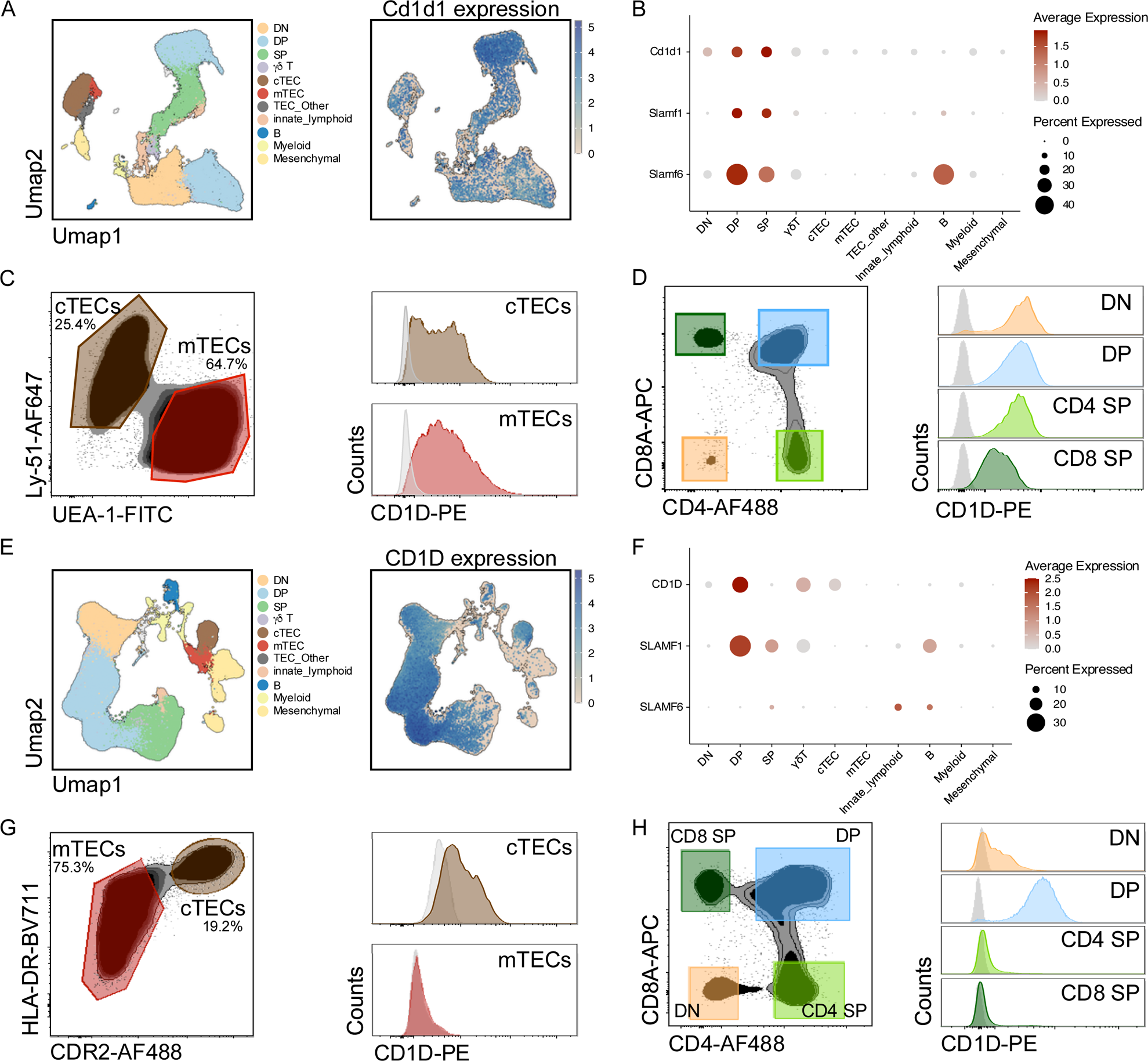
CD1D gene and protein expression in mouse and human thymus. A/E: Clustering of thymic cell populations and their expression of Cd1d1 (mouse)/CD1D (human) derived from the mouse and human thymus cell atlas, respectively (Ref (Park *et al*., 2020)). B/F: Normalized expression of Cd1d1/CD1D and Slam/SLAM transcripts across thymic cell populations. Flow cytometry of mouse and human TECs (C/G) and thymocyte subsets (D/H).

## Discussion

In this study, we employed multi-modal single-cell transcriptomics to explore the diverse phenotypic states that T_inn_ cells can manifest within the human thymus and blood. Through a comprehensive analysis, we juxtaposed these states with those of conventional T cells, providing novel insights into human T cell biology and a comprehensive resource for further studies of health and disease. T_inn_ cells have garnered substantial attention recently because of their distinctive developmental pathway and functional characteristics, which are increasingly being explored for potential applications in immunotherapies (Delfanti et al., 2022; Dogan et al., 2022; Lee et al., 2023).

Our study demonstrated that in human blood, the majority of T_inn_ cells exhibit a distinct transcriptional program that is shared by most iNKT, MAIT, and Vδ2Vγ9 T cells under steady-state conditions. This program implies a blended type I/type III transcriptional pattern, driven by specific transcription factors that enable the expression of distinct chemokine and cytokine receptors, NK receptors, and cytotoxic molecules. This program equips T_inn_ cells with the ability to swiftly respond to cytokines like IL-12, IL-18, and IL-23, independently of TCR signaling (Philippot et al., 2023; Ussher *et al*., 2014). Notably, we and others (Duquette *et al*., 2023; Kurioka et al., 2015) showed that human T_inn_ cells constitutively express granzyme K but lack granzyme B, while also expressing cathepsins, which are necessary for activating granzymes (D’Angelo et al., 2010). This indicates that T_inn_ cells are poised to release active granzyme K upon stimulation (Kurioka *et al*., 2015). Granzyme K possesses a range of immunomodulatory functions. It can induce the production of pro-inflammatory cytokines such as IL-6 and IL-8 from epithelial cells (Kaiserman et al., 2022) and of IL-6, CCL5, and CCL2 from fibroblasts (Jonsson *et al*., 2022). Mouse granzyme K can trigger the maturation and secretion of pro-inflammatory interleukin-1b, particularly in LPS-sensitized peritoneal macrophages (Wensink et al., 2014). Additionally, granzyme K may activate a novel complement pathway independently of the classical, alternative, and lectin pathways (Jonsson et al., 2023), implying its involvement in immune regulation and inflammatory responses. Given this overarching role of granzyme K in immune regulation, it appears that the initial role of human T_inn_ cells upon activation may indeed be the release of granzyme K, which likely happens concomitantly or before cytokine secretion or cytotoxic activity. In contrast, mouse T_inn_ cells do not express granzyme K transcripts but unlike human T_inn_ cells already possess pre-formed cytokine-encoding transcripts even before any stimulation occurs, allowing for an immediate response (Govindarajan et al., 2018; Matsuda et al., 2003). This suggests that despite their evolutionary conservation, T_inn_ cells may have evolved species-specific mechanisms to provide early signaling and amplification of the adaptive immune response.

We identified a set of transcription factors and their predicted target genes that exhibited increased transcriptional activity in human T_inn_ cells when compared to naive and effector T_conv_ cells. Several of these transcription factors have previously been associated with the development and function of mouse T_inn_ cells, and we found that the expression of some of them is shared between species. Notably, several of these transcription factors are associated with the differentiation and production of IFNγ, cytotoxicity (Intlekofer *et al*., 2005; Istaces *et al*., 2019; Klein-Hessling *et al*., 2017; Nah and Seong, 2022) as well as the production of the cytokine IL-17 (Chang *et al*., 2019; Ciofani *et al*., 2012; Yu *et al*., 2017). These connections are consistent with the type I/type III transcriptional program observed in T_inn_ cells. In mice, the well-established Th1/Th17 paradigm identifies IL-12 as a cytokine that induces IFNγ production, while IL-23 is known for inducing IL-17 production. This is in line with the varied expression of IL-12R and IL-23R receptors observed on mouse type I and type III T_inn_ cell subsets, which correlates with their respective cytokine production profiles. By contrast activating human MAIT cells either through their TCR or with a combination of IL-12 and IL-18, as well as stimulating T_inn_ cells with IL-23, all result in the production of IFNγ (Garner *et al*., 2023; Philippot *et al*., 2023). Yet, only a subset of these cells produces IL-17 in the same conditions, a phenomenon thought to be influenced by epigenetic modifications at the IL-17 gene loci (Garner *et al*., 2023). Underlining the importance of understanding the regulation of IL-17 production in T_inn_ cells, there has been an increase in IL-17 production observed in human MAIT cells in cases including severe asthma, community-acquired pneumonia in children (Lezmi et al., 2019; Lu et al., 2020) and colorectal cancer patients (Borras et al., 2023). Interestingly, we found NR1D family transcription factors, which have been previously associated with regulation of Th17 cells (Chang *et al*., 2019; Yu et al., 2013), are driving regulons in human T_inn_ cells. NR1D TFs are regulated by the body’s circadian clock, suggesting that the circadian rhythm might affect how T_inn_ cells produce IL-17, a hypothesis that warrants further investigation. While many of the TFs essential for establishing the human T_inn_ program were also expressed in mouse T_inn_ cells, there were notable exceptions. For example, CEBPD, EOMES, and MYBL1 were found to be highly expressed in human T_inn_ cells under steady-state conditions, yet we did not detect transcripts for these TFs in the mouse T_inn_ reference dataset. CEBPD has previously been implicated in regulating the expression of CCR6 in human MAIT cells(Lee *et al*., 2018). However, its predicted targets, including ZBTB16, suggest that it may play a crucial role in regulating the human T_inn_ program. MYBL1’s preferential expression in T_inn_ cells has been previously observed (Gutierrez-Arcelus et al., 2019), but its specific function in these cells remains to be defined. EOMES has been reported to play a role in the development of mouse iNKT cells, although its expression level is very low under steady-state conditions (Shimizu et al., 2019; Townsend et al., 2004). By contrast, Tbet is highly expressed in type I mouse T_inn_ cells and is essential for their development and functions (Matsuda et al., 2006; Townsend *et al*., 2004). However, human effector T_inn_ cells, which are most similar to mouse type I T_inn_ cells, express relatively low levels of Tbet. Instead, Tbet’s expression and activity were correlated with the acquisition of the GEP6 program by T_conv_ cells in humans. These findings suggest the possibility of species-specific transcriptional regulation of T_inn_ cells, which could be relevant for future therapeutic applications involving these cells. Curiously, high confidence regulons like PLZF and Rorγt were not identified in the gene regulatory network of human T_inn_ cells, possibly due to the relatively low gene detection in this context.

In the post-natal thymus, we observed iNKT and MAIT cells that displayed a transcriptional profile similar to developing conventional CD4^+^ and CD8^+^ T cells, respectively. This finding raises several possible interpretations. One plausible scenario is that these naïve T_inn_ cells could potentially serve as precursors to effector T_inn_ cells. This implies that T_inn_ cells likely undergo a maturation process, which could occur either within the thymus or in peripheral tissues, facilitating the acquisition of effector functions. Notably, in human cord blood, iNKT and MAIT cells with a naïve phenotype are more prevalent, and there is a gradual increase in the proportion of effector T_inn_ cells with age. If this hypothesis is accurate, it would suggest that T_inn_ cells initially experience positive selection in a manner akin to conventional T_conv_ cells or that their mode of selection might not provide the necessary signals for complete maturation. They would then subsequently receive distinct signals that would propel them to acquire effector functionalities. This concept finds support in recent studies involving Vγ9Vδ2^+^ T cells. In this context, immature naïve-like CD4^+^ CD161^−^ cells were observed to undergo a transition toward an effector transcriptome (Perriman *et al*., 2023). During this transition, there was an upregulation of genes encoding various cytotoxic molecules, chemokines, chemokine receptors, as well as different cell surface markers (Perriman *et al*., 2023). This maturation could be recapitulated *in vitro* by culturing naïve Vγ9Vδ2^+^ T cells with OP-9 cells in the presence of IL-2, IL-7, and IL-15 cytokines. Our analysis of human thymic γδ T cells highlighted the presence of transcriptionally transitional cells, reinforcing the notion of a sequential developmental trajectory. Although this scenario could potentially extend to human iNKT and MAIT cells, another intriguing possibility could be envisioned. In addition to the pool of naïve T_inn_ cells, a minority of cells in the human thymus expressed genes linked to effector functions or genes typically associated with T_inn_ cells. Notably, these effector cells formed distinct transcriptional clusters and despite analysis of several thousand cells, we did not identify transitional cells bridging the gap between the naïve and effector populations. This leads us to speculate that naïve thymic T_inn_ cells may not necessarily serve as precursors to effector T_inn_ cells. Instead, these distinct populations could potentially represent the results of two separate selection pathways for T_inn_ cells. Supporting this idea, naïve MAIT cells (and some naïve iNKT cells) have been identified in the mouse thymus (Chandra *et al*., 2023; Krovi *et al*., 2022; Legoux *et al*., 2019). These cells are thought to be T_inn_ cells that have undergone positive selection on TECs, which do not express the SLAM family receptors necessary for the acquisition of the effector program. Indeed, we showed that both human and mouse cTECs express CD1d molecules on their cell surface. Interestingly, we also observed species-specific differences, notably the lack of CD1d expression in human mTECs compared to mouse mTECs. This distinction could contribute to variations in the maturation of iNKT cells between humans and mice. In the absence of IL-15 cross-presentation by mTECs, mouse iNKT cells do not fully mature (Cui *et al*., 2022; White *et al*., 2014). CD1d crosslinking has been shown to stimulate the production of IL-15 in epithelial cells from the reproductive tracts (Kawana et al., 2008). Taken together, the absence of CD1d on human mTECs might explain why mouse iNKT cells can mature when they interact with CD1d expressed on mTECs, while this maturation process cannot occur in humans. These findings underscore the intricate nature of T_inn_ cell development and maturation, suggesting the existence of multiple potential pathways and mechanisms, some of which may be species-specific and require further experimental investigation.

Finally, our study highlights a distinct path taken by T_inn_ cells with an effector program in the human postnatal thymus, characterized by a mixed type I/type III effector potential. This contrasts with mice, where T_inn_ cells tend to split into multiple effector subsets. Interestingly, we observed a cluster of proliferating cells among human γδ T cells in the thymus, but a similar proliferative cluster was absent for human iNKT and MAIT cells. This also contrasts with mice, where clusters of thymic iNKT and MAIT cells undergoing cell division are readily identifiable (Baranek *et al*., 2020; Harsha Krovi *et al*., 2020; Legoux *et al*., 2019), reflecting the proliferative burst following positive selection(Benlagha et al., 2002), crucial for establishing a substantial pool of T_inn_ cells. Consequently, while T_inn_ cells constitute around 1-2% of thymocytes in mice, their proportion is at least one order of magnitude lower in pediatric humans. Moreover, our analysis did not reveal any T_inn_ cells with a type II transcriptome in humans, unlike in mice. In mice, thymus-resident iNKT2 cells serve as a major source of IL-4 (Lee *et al*., 2013), significantly impacting the thymic environment. This IL-4 influence includes effects on thymocyte emigration(White et al., 2017), induction of memory-like traits in CD8^+^ T cells (Lee *et al*., 2013; Weinreich et al., 2010), and activation of specific dendritic cells to produce chemokines that promote clonal deletion, all while sparing regulatory T cells (Breed *et al*., 2022). Additionally, iNKT2 cells in mice contribute to the formation of the thymus medulla through RANK signaling (White *et al*., 2014). The scarcity of type II T_inn_ cells in the human thymus suggests that these phenomena may be species-specific or regulated by different cell types in humans.

Taken together, our findings hold significance in elucidating the diverse functional attributes of T_inn_ cells and their potential applications in immunotherapeutic contexts.

## Material and Methods

### Mice

The *Cd1d1d2*^−/−^ mice backcrossed to the C57BL/6 background have been described previously(Chen et al., 1997). C57BL/6 were purchased from Jackson Laboratories. All mice were used between 6 to 15 weeks and were age matched for each experiment. Mice were raised in a specific pathogen-free environment at the Office of Laboratory Animal Research at the University of Colorado Anschutz Medical campus or the Animal Core Facility at Cold Spring Harbor Laboratory. Animal procedures were approved by the UCD (00065) Institutional Animal Care and Use Committees and the Cold Spring Harbor Laboratory IACUC (23-1); all procedures were carried out in accordance with the approved guidelines.

### Mouse samples

To isolate thymocytes, thymus tissue was immersed in RPMI 1640 media (Corning, #10-040-CV) and gently pressed through a 40μm cell strainer using the plunger of a 1 mL syringe. For TEC isolation, the thymus tissue was cut into small fragments and submerged in RPMI 1640 media without phenol red (Gibco, #11835030), supplemented with 20mM HEPES (Gibco, #15630080), 1.3 U/mL Liberase TH (Sigma-Aldrich #5401135001), and 100 U/mL DNase I (Roche, #11284932001). These tissue fragments were incubated for 5 minutes on ice followed by an additional 20 minutes at 37°C. After the digestion period, the solution was repeatedly mixed with a micropipette to ensure complete tissue disintegration. To stop the digestion process, cells were suspended in HBSS, 4% heat-inactivated FBS (HI-FBS, FBS from Corning, #35-010-CV, preheated for 20 minutes at 56°C), 20mM HEPES, and 10U/mL DNase I. To remove immune cells, the cells were incubated with rat anti-mouse CD90.2 (clone 53-2.1, Biolegend #140302), anti-mouse CD45 (clone 30-F11, Invitrogen #14-0451-85), and anti-mouse CD45-BV605 (clone 30-F11, Biolegend #103139) antibodies for 30 minutes at 4°C. Subsequently, the cells were placed on panning plates coated with goat anti-rat IgG (Vector Laboratories, #BA-9400) for 20 minutes at room temperature. Unattached cells were then transferred to new panning plates for a second round of depletion. The remaining cells, following this depletion process, were prepared for flow cytometry analysis.

### Human samples

De-identified Peripheral blood samples from healthy donors were obtained through the University of Colorado Clinical and Translation Research Centers (CTRC), which is a part of the Colorado Clinical and Translation Sciences Institute (CCTSI). These samples were collected using sodium heparin tubes, and peripheral blood mononuclear cells (PBMCs) were isolated using a Ficoll gradient provided by Cytiva. Additional samples were acquired from plateletpheresis leukoreduction filter chambers (LRS) obtained from the Vitalant Blood Center located in Denver, Colorado, USA. For pediatric thymus tissues, which were extracted from infants undergoing corrective surgeries for congenital heart disease, the processing was initiated within one hour of extraction. These tissue samples were procured from various sources, including Children’s Hospital Colorado, the Mount Sinai, and the Northwell Health Biorepository, following ethical approval (IRB 20-0150, NHBR2101). Pediatric thymus samples for scRNAseq came from individuals between 10 and 20 weeks old (Supplementary Table 1), and samples used for flow cytometry experiments came from individuals between 4 days and 5 months old. To extract thymocytes for both single-cell RNA sequencing (scRNAseq) and flow cytometry, the thymus tissue was placed in complete RPMI 1640 media (Gibco, #22400-071) (10% heat-inactivated fetal bovine serum (FBS, Sigma-Aldrich), 1% non-essential amino acids (Sigma-Aldrich), 1% Sodium Pyruvate (Sigma-Aldrich), 1X GlutaMAX (Gibco), 1% Penicillin/Streptomycin (Gibco), and 1X 2-mercaptoethanol (BME, Sigma-Aldrich)), cut into small pieces, and gently pressed with the back of a 10 ml syringe to release thymocytes. The resulting suspension was passed through a 70 µm filter. Thymocytes and PBMCs were isolated using a Ficoll-Paque density gradient provided by Cytiva. PBMCs were cryopreserved in FBS with 10% DMSO from Sigma-Aldrich and stored in liquid nitrogen. Tetramer staining for MAIT and iNKT cells was performed on freshly isolated thymocytes. For tetramer staining of MAIT and iNKT cells, freshly isolated thymocytes were used. To enrich TECs for flow cytometry, thymus tissue was cut into small pieces and placed in RPMI 1640 media without phenol red, 5% heat-inactivated FBS, 1% Penicillin/Streptomycin, 10mM HEPES (Gibco, #15630080), and 0.55mM 2-mercaptoethanol (Gibco, #21985023). The thymus tissue in this media was stirred on a magnetic plate for 40 minutes. The supernatant was removed and replaced with fresh media every 10 minutes to remove released thymocytes. The remaining tissue chunks were placed in a digestion buffer consisting of RPMI 1640 media without phenol red, 2% HI-FBS, 20mM HEPES, 80 U/mL DNase I (Roche, #11284932001), 1.6 U/mL Dispase I (Roche, #04942086001), and 0.3mg/mL Collagenase IV (StemCell Technologies, #07427) for digestion at 37°C with gentle shaking. This digestion process was conducted in two sessions of 25 minutes each, with the supernatant being extracted and replaced with fresh digestion buffer in between. At the end of the digestion, the tissue chunks had nearly entirely disintegrated, and the digestion was halted by resuspending cells in the same buffer used for thymocyte release. The combined supernatants were further incubated in TrypLE Express Enzyme (Gibco, #12604-013), 1mM MgCl_2_, 2mM CaCl_2_, 100U/mL DNase I for 5 minutes at 37°C to obtain a single-cell suspension. The digestion was stopped by resuspending cells in the previously described buffer. To remove immune cells and erythrocytes, cells were incubated with mouse anti-human CD3 (clone UCHT1, Biolegend #300402), anti-human CD4 (clone RPA-T4, Biolegend #300570), anti-human CD8 (clone RPA-T8, Biolegend #301002), anti-human CD45 (clone HI30, Biolegend #304002) and anti-human CD235a (clone HI264, Biolegend #349102) antibodies in HBSS (Gibco, #14175079), 4%HI-FBS and 20U/mL DNase I, for 30mins at 4°C. Cells were then placed on panning plates coated with goat anti-mouse IgG (Vector Laboratories, #AI-9200-1.5) for 20 minutes at room temperature, and the unadhered cells were transferred to new panning plates for a second round of depletion. The remaining cells following depletion were then stained for flow cytometry. Overview of sample metadata is provided in Supplementary Table I.

### Magnetic-bead enrichment of iNKT and MAIT cells

To enrich for thymic MAIT and thymic/blood iNKT cells, up to 2×10^9^ cells were incubated with MR1-5-OP-RU-PE-Tet or CD1d-PBS57-PE respectively in MACS buffer (0.5% BSA, 2mM ETDA, PBS), for 25 mins at room temperature. Cells were washed twice and incubated with anti-PE microbeads (Miltenyi), followed by separation using an autoMACS Pro Separator (Miltenyi) according to manufacturer’s instructions. PE-microbead-labelled cells in the enriched fraction were stained with the specified panel of antibodies listed below.

### Fluorescence-activated cell sorting

Single cell suspensions were stained with efluor780 viability dye (ThermoFisher) for 10 mins at room temperature and washed once prior to cell surface staining. Enriched MAIT and iNKT from thymus, enriched iNKT from PBMC, unenriched γδ T, CD4^+^ and CD8^+^ from thymus, and unenriched MAIT, γδ T, CD4^+^ and CD8^+^ T cells from PBMC were stained with the following cell surface markers in MACS buffer at room temperature for 20 mins: CD3-AF488 (clone OKT3, Biolegend), CD14-eFluor450 (clone 61D3, ThermoFisher), CD19-eFluor450 (clone H1B19, ThermoFisher), Vα7.2-BV785 (clone 3C10, Biolegend), Vα24-PerCP-Cy5.5 (clone C15, Biolegend), CD4-AF710 (clone OKT4, Tonbo), CD8α-PE-Cy7 (clone SK1, Tonbo), TCRγδ-BV650 (clone 11F2, BD Biosciences), FcγR block (Miltenyi). Cells were washed twice and resuspended in MACS buffer prior to cell sorting on the Aria 3 (BD Biosciences). Purified cell populations were sorted into MACS buffer. To confirm gene expression from scRNAseq analysis, MAIT and iNKT cells were enriched from the human thymus as described above, as were iNKT cells from human blood. γδ T cells were stained directly from the human thymus as were blood MAIT and γδ T cells. Single cell suspensions were stained as above with efluor780 viability dye prior to incubation at 37C for 10 min with CCR7-APC-Fire810 (clone G043H7, Biolegend) and FcγR block. A combination of the following cell surface markers were subsequently added and cells were stained at room temperature for 15 min: CD3-BUV496 (clone UCHT1, BD Biosciences), CD14-PE-Cy5 (clone 61D3, ThermoFisher), CD19-PE Cy5 (clone H1B19, ThermoFisher), Vα7.2-BV785 (clone 3C10, Biolegend), Vα24-PerCP-Cy5.5 9clone C15, Biolegend), CD4-BV570 (clone RPA-T4, Biolegend), CD8α-BUV395 (clone RPA-T8, BD Biosciences), TCRγδ-BV650 (clone 11F2, BD Biosciences), Vδ1-PerCP-Vio700 (clone REA173, Miltenyi), Vδ2-FITC (clone 123R3, Miltenyi), Vγ9-PE (clone B3, BD Biosciences), CD161-BUV805 (clone HP-3G10, BD Biosciences), CD62L-BV650 (clone DREG-56, Biolegend). Cells were then washed twice with MACS buffer prior to fixation and intracellular staining performed with BD Transcription Factor Buffer Set according to the manufacturer’s specification. The following antibodies were used to stain for intracellular proteins: PLZF_PE-CF594 (clone R17-809), Eomes-BUV737 (clone X4-83), Tbet-BV605 (clone 4B10, Biolegend), GZMK-AF660 (clone G3H69, InVitrogen), GZMB-AF700 (clone GB11, BD Biosciences). Phenotypic analyses and validation of the cell sorting panel was performed on the Cytek Aurora flow cytometry system using SpectroFlo software (v3.0). Data were analyzed using FlowJo software v10.7.1 (BD Biosciences).

### Flow cytometry analysis of CD1d expression

For the mouse experiments, thymocytes were resuspended in PBS, 5% FBS (Corning, #35-010-CV), 4mM EDTA and stained for 30mins at 4°C with: Fc blocker CD16/32 (clone 93, Invitrogen #14-0161-85), CD4-AF488 (clone GK1.5, Biolegend #100423), CD8α-APC (clone 53-6.7, Biolegend #100711), CD1d-PE (clone 1B1, Biolegend #123509). For the murine thymus samples that were depleted of immune cells, the single cell suspension was resuspended in HBSS, 4% HI-FBS, 20mM HEPES, 10U/mL DNaseI, 2.5mM EDTA, and stained for 30mins at 4°C with: Fc blocker CD16/32 (clone 93, Invitrogen #14-0161-85), Epcam-BV421 (clone G8.8, Biolegend #118225), CD45-BV605 (clone 30-F11, Biolegend #103139), UEA1-FITC (Vector Laboratories, #FL-1061-2), Ly-51-AF647 (clone 6C3, Biolegend #108312), and CD1d-PE (clone 1B1, Biolegend #123509). For the flow cytometry experiments on human samples, thymocytes were resuspended in PBS, 2% FBS, and stained for 30mins at 4°C with: TruStain FcX (Biolegend #422302), CD45-BV421 (clone HI30, Biolegend #304032), CD4-AF488 (clone OKT4, Invitrogen #53-0048-42), CD8-APC (clone RPA-T8, Invitrogen #17-0088-42), CD1d-PE (clone 51.1, Biolegend #350305). For the human samples that were depleted of immune cells and erythrocytes, cells were resuspended in PBS, 2% FBS, and stained for 30mins at 4°C with: TruStain FcX (Biolegend #422302), CD45-AF647 (clone QA17A19, Biolegend #393406), EPCAM-BV421 (clone 9C4, Biolegend #324220), CDR2-AF488 (pure CDR2 antibody kindly provided by Dr. Sheena Pinto, conjugated with the AF488 antibody labeling kit by Invitrogen, #A20181), HLADR-BV711 (clone L243, Biolegend #307643), CD1d-PE (clone 51.1, Biolegend #350305). In all experiments, to measure viability cells were stained with the live/dead Fixable Near-R dead cell stain kit (Invitrogen #L10119). Flow cytometry was performed on a BD LSR Fortessa Cell Analyzer.

### Single-cell RNA-sequencing

Single cell whole transcriptomes and TCR sequencing libraries were prepared using the BD Rhapsody Single-Cell Analysis System (BD Biosciences) according to the manufacturer’s specifications. Prior to cell sorting on the Aria 3 (BD Biosciences) and during cell surface antibody staining, up to 2 x10^6^ enriched or unenriched cells were labeled with an oligonucleotide-tagged antibody sample tag (BD Biosciences). From infant thymus and PBMC donors up to 5 populations were sorted after doublet, viability, B cell (CD19^+^CD3^−^) and monocyte (CD14^+^CD3^−^) discrimination: 1. MAIT cells (MR1-5-OP-RU-Tet^+^Vα7.2^+^CD3^+^), 2. iNKT cells (CD1d-PBS57-Tet^+^Vα24^+^CD3^+^), 3. γδ T cells (CD3^+^TCRγδ^+^), 4. CD4^+^ T cells (CD4^+^CD8^−^CD3^+^) and CD8^+^ T cells (CD8^+^CD4^−^CD3^+^). Cell subsets sorted for the different donors are listed in Supplementary Table I and the gating strategy is shown in Supplementary Fig. 15. Prior to cDNA library preparation for the WTA and VDJ libraries, all cell subsets from the different donors were pooled, with up to 12 unique sample tags combined per library. Libraries were quantified and pooled according to equivalent molar concentrations and sequenced on the NovaSeq sequencing platform at the University of Colorado Genomics Core with the following read lengths: read 1 – 150 cycles; read 2 – 150 cycles; and i7 index - 8 cycles.

### Single-cell RNA-seq data analysis

The quality of sequencing reads was evaluated using FastQC and MultiQC. Sequencing reads (FASTQ) were mapped and sample Tag deconvoluted with The BD Rhapsody™ WTA Analysis Pipeline on the GRCh38 genome sequence. This pipeline produced a gene expression matrix for each sample, which records the number of UMIs for each gene associated with each cell barcode. Aggregated data were then imported into the R environment and analyzed with Seurat (4.3.0). Low-quality cells were filtered using the cutoffs nFeature_RNA >= 500 & nFeature_RNA < 3000. Cells with a high mitochondrial content were removed using a batch-dependent threshold with the isOutlier function from the Scater package (McCarthy et al., 2017). Genes expressed in less than 20 cells were ignored. This resulted in 78,607 cells with 17,204 genes for downstream analyses. The NormalizeData function of Seurat was performed using default parameters to remove the differences in sequencing depth across cells. Dimensionality reduction was performed prior to integration for visualization purposes (Supplementary Fig 2A), by selecting 2000 highly variable genes for principal component analysis (PCA) and uniform manifold approximation and projection (UMAP). To integrate the data and remove batch-effects from the PCA subspaces based on the correct cell alignment, we used Harmony (Korsunsky et al., 2019) following PCA to project cells into a shared embedding in which cells group by cell type rather than dataset-specific conditions. We then applied the RunUMAP function on 20 dimensions of the harmony embedding to obtain bidimensional coordinates for each cell. We determined the k-nearest neighbors of each cell using the FindNeighbors function and used this knn graph to cluster cells using the Louvain algorithm from FindClusters based on the same harmony dimensions as the RunUMAP function (20 dimensions, resolution 1.2). This dataset was subsequently split up into 5 cell types from 2 different tissues based on cell hashing tags/barcodes. Each cell type from each tissue was re-analyzed individually using the same steps to obtain UMAPs and clusters in Figures 2 and 4. Plots displaying cells on UMAPs were generated using the SCpubR package (v1.1.2) (Blanco-Carmona, 2022).

### LISI metric and analysis of cluster stability

The local inverse Simpson’s index (LISI) was used to assess the degree of mixing during batch correction and dataset integration in scRNA-seq analysis (Korsunsky *et al*., 2019). This approach helps evaluate the effectiveness of data integration methods by quantifying how well datasets are merged without introducing artificial batch effects. To assess the integration process, we employed the integration LISI (iLISI) score. iLISI measures the effective number of datasets within a neighborhood and provides an indication of how effectively the individual datasets have been harmoniously integrated into a unified whole during the analysis. In addition, we used the “cell-type” LISI (cLISI) score to evaluate the accuracy of cell-type assignments in the integrated dataset. cLISI is a modified version of the LISI score, but instead of assessing dataset labels, it focuses on the accuracy of cell type assignments within the integrated data. As the specific identities of individual cells were not known beforehand, we assigned mock cell identities based on anticipated gene expression patterns. These mock identities were determined using prior knowledge of gene expression markers associated with distinct cell types. For instance, we identified DN thymocytes as cells expressing PTCRA > 1, B cells as cells expressing CD19 > 1 and IGKC > 1, T_regs_ as cells expressing FOXP3 > 1, MAIT cells as cells expressing SLC4A10 > 1 and FOXP3 < 1, CD4 T cells as cells expressing CD4 > 1, CD8A < 1, SLC4A10 < 1, FOXP3 < 1, and CCR7 > 1, DP thymocytes as cells expressing RAG1 > 1 and CD1C > 1, and CD8αα thymocytes as cells expressing CD8A > 1 and GNG4 > 1. These mock identities were used as initial cell type assignments and served as the basis for assessing the success of integration, as indicated by increased iLISI scores and the maintenance of a cLISI score of 1. Only cells with assigned mock identities were included in the cLISI analysis. To evaluate the stability of clusters, we conducted a bootstrapping procedure in which cells from each predefined cluster were repeatedly sampled and then subjected to re-clustering. Cluster stability was assessed by examining co-assignment probabilities (CP), where higher CP values indicated greater cluster stability. In essence, a high CP suggests that the cells within a cluster consistently grouped together across multiple iterations, reinforcing the reliability and robustness of that cluster’s identity.

### TCR analysis

V(D)J single cell sequencing data were mapped and quantified using the BD Rhapsody™ WTA Analysis Pipeline and the GRCh38 genome sequence. To connect the VDJ data with transcripts data for each cell, we established links based on cell indexes extracted from the consensus annotation files (VDJ_percell.csv) and MolsPerCell.csv files from each demultiplexed sample. Only TCR paired sequences were retained for subsequent analyses. TCR data from each VDJ-sequenced sample were combined together and added to the metadata of the Seurat object. Clonotypes were defined based on unique TCR VJ usage and complementary-determining region (CDR3) motifs. Basic TCR statistics, such as the distribution of length and counts were computed using the tidyverse package (v1.3.2). The assessment of clonotype diversity was conducted using the mean value of the Shannon index, computed through the diversity function of the vegan R package (v2.6-4) after 100 iterations. Prior to the diversity calculation, the data was subjected to rarefaction to match the lowest sequence count found within the studied groups. Chord diagrams were generated using the circlize package (v0.4.15) (Gu et al., 2014) and CDR3 motif logos using the ggseqlogo package (v0.1) (Wagih, 2017). The stacked letters’ cumulative height at each position signifies the degree of sequence conservation, portraying the relative abundance of amino acids, which is further depicted by the varying heights of individual letters within the stack.

### Identification of differentially expressed genes between clusters

We identified cluster-enriched genes by using the FindAllMarkers function in Seurat with test.use = wilcox. This function identified differentially expressed genes for each cluster by comparing the gene expression for cells belonging to a cluster versus cells belonging to all other clusters. Only those genes that passed an adjusted *p* value (Benjamini-Hochberg) cutoff of 0.05, log fold change > 0.4 and min.pct = 0.3 were included in the downstream analyses.

### Characterizing the replicability of cell types defined by scRNA-seq between studies and between species

We assessed the consistency of cell clusters in our integrated thymic data by comparing them with the human thymus atlas from the Park *et al*. dataset (Park *et al*., 2020). To do this, we focused exclusively on thymocytes, totaling 37,369 cells in our dataset. We also acquired the annotated AnnData object from the Park *et al*. dataset, which specifically contained T cells. To enable a meaningful comparison, we combined the two raw count matrices, concentrating on the top 2000 highly variable genes shared across both datasets. This resulted in a matrix containing 3,106 genes and 114,363 cells. To evaluate the consistency of cell types between these datasets, we employed the pyMN package to perform unsupervised MetaNeighbor analysis (Crow *et al*., 2018). MetaNeighbor assesses the similarity of cell types by constructing a network of cells based on the correlation of their gene expression profiles. It then predicts cell type labels, hiding them from one dataset while using the other. The result is expressed as a mean Area Under the Receiver Operator Characteristic (AUROC) score, which measures the probability of correctly identifying a cell’s type based on its gene expression profile. We used the ggplot2 package to visualize the AUROC scores obtained from pyMN, comparing our integrated clusters with the thymocyte clusters defined in the Park *et al*. dataset. For assessing the replicability of cell clusters across species, we utilized the reference scRNAseq murine T_inn_ dataset and our human thymic iNKT, MAIT, and γδT individual seurat objects from Figure 2. To ensure an appropriate comparison, we obtained orthologous genes between mouse and human using the biomaRt package (Durinck et al., 2005; Durinck et al., 2009). We filtered the murine count matrix to retain only genes with known 1:1 orthologs in humans. Then, we performed unsupervised MetaNeighbor analysis with pyMN on the combined set of highly variable genes from both human and mouse datasets. Finally, we used ggplot2 to create visualizations of the AUROC scores returned by pyMN, including clusters that contained at least 1% of the cells in each species to ensure greater confidence in assessing the replicability of clusters across species.

### Identification of Gene Expression Programs

The count matrix was used for conducting non-negative matrix factorization (NMF) through the cNMF method (Kotliar *et al*., 2019). This process enabled us to infer both identity and activity programs, along with their respective contributions in each cell. The usage of each program for each cell was added to the metadata of the Seurat object and displayed as a featurePlot. To determine the genes associated with each program, we plotted the gene ranks (ranging from most associated to least associated) against the gene_spectra_score output from the cNMF analysis. The plotted gene ranks were fitted to a sigmoid curve and the slope at the first elbow point was calculated as the minimum threshold for genes to be retained in a given GEP. The same slope was applied to every GEP to prevent bias in ranked gene selection, as the gene rankings between GEPs are not comparable and are relative to each GEP (as depicted in Supplementary Figure 16). Cells from the blood sample were assigned to the GEP with the highest usage (as provided by cNMF), to display an alluvial plot with ggalluvial (v0.12.5) (Brunson, 2020).

### Scoring of Gene Signatures

Gene signatures were scored on our Seurat object, or on other dataset’s Seurat or AnnData objects using either the function AddModuleScore in Seurat, or scanpy.tl.score_genes in scanpy. In both cases, the score is computed as the average expression of all genes contained in the gene list, and subtracting the average expression of 100 control genes (randomly chosen to match the expression bins of the gene list). Gene signatures used throughout this manuscript and their source can be found in supplementary Table 2.

### Gene regulatory network inference

To deduce gene regulatory networks, we employed pySCENIC from a pre-built singularity container, aertslab/pyscenic:0.12.1, a tool utilizing cis-regulatory motif analysis to identify potential transcription factors (TFs) that might govern a cluster of co-expressed genes within individual cells (Aibar *et al*., 2017). pySCENIC was run using the – mask-dropouts flag and a normalized enrichment score threshold of 2 to help mitigate the effects of the varying degrees of sparsity across the data sets we generated. The initial step involved generating modules composed of transcription factors and co-expressed genes using GRNboost2 (Ref (Moerman et al., 2019)). These modules were pruned to remove indirect targets that lacked significant enrichment for the corresponding TF motif within ±10 Kb from the transcription starting site of the putative target (cisTarget). This process yielded a collection of transcription factor regulons. Considering the inherent stochasticity in gene regulatory network inference using GRNBoost2, each run of pySCENIC may yield different quantities of regulons, along with distinct target genes associated with each TF. To mitigate this variability, we performed 100 pySCENIC runs and retained regulons present in 100% of the runs. We also removed regulons that did not have at least 5 target genes defining the regulon activity. Due to the high degree of noise in target genes, we retained target genes that appeared within a regulon in at least 95% of the runs. Furthermore, each target gene also had to overlap with the union of all possible retained ranked gene expression targets across all GEPs generated from cNMF. To identify regulons that were specific to the underlying biology of our cell types and GEPs, we calculated the AUC scores using the R package AUCell, located in the pySCENIC container, for each regulon based on the pruned target gene list. A regulon was deemed specific to a defined cell population if at least 20% of the cells within the annotated population scored in the 90^th^ percentile of the overall AUC score for all cells.

### Comparison of Gene Expression Programs with gene signatures from the literature

We obtained gene signatures identified from (1) differential expression (DE) analysis from bulk RNAseq between sorted naïve, T_cm_, T_em_ CD4^+^ and CD8^+^ T cell populations by Rose *et al*. (Rose *et al*., 2023); (2) DE genes between cell clusters defined from scRNAseq of naïve and memory CD4^+^ T cells isolated from PBMCs by Cano-Gamez *et al*. (Cano-Gamez *et al*., 2020); (3) DE genes between cell clusters defined from scRNAseq of blood immune cells by Terekhova *et al*. (Terekhova *et al*., 2023) (4) DE genes between cell clusters defined from scRNAseq of T cells across nine human tissues by Poon *et al*. (Poon *et al*., 2023). In the Rose dataset, we kept genes that defined their Figure 2 E, H (adjusted p-value ≤ 0.05). In the Cano-Gamez and Poon datasets, we kept DE genes with a minimum log fold-change of 0.25 (adjusted p-value threshold ≤ 0.05 or 0.01, respectively). In the Terekhova dataset, we used the top 100 differentially expressed genes shared in their supplementary Table S5. We computed the Jaccard Index (JI) between the gene lists derived from our GEPs and those from the Rose, Cano-Gamez, Terekhova and Poon datasets. Since the gene lists varied in length, we weighted the JI to make it comparable across pairwise comparisons. This was achieved by dividing the JI by the maximal theoretical JI for each pairwise comparison, which is the ratio of the length of the smaller list to the length of the larger list. To assess the significance of the observed JI, we performed a permutation analysis. We generated 1000 random gene lists A’ and B’, matching in length and expression pattern to the original lists A and B. We computed the weighted JI between these random lists and defined an empirical p-value by counting how many of these weighted JIs were greater than the observed weighted JI divided by the number of permutations. To account for multiple comparisons, we applied a Bonferroni correction to the empirical p-values. For the co-expression analysis of GEPs and gene lists from other datasets, we scored the gene lists on the entire integrated dataset. This was done using functions like Seurat’s AddModuleScore with the blend=TRUE parameter. Additionally, GEP4, GEP5, and GEP6 were scored on the Poon et al. dataset using scanpy’s tl.score_genes function, and their scores in specific cell clusters of interest were displayed in Supplementary Figure 9B.

### Pseudo-bulk differential expression analysis

To investigate for cell lineage-specific gene signatures in PBMCs, we grouped cells by batch, cluster and lineage, restricting our analysis to only batches E, F and I where at least 3 or more cell lineages were sorted and sequenced within the same batch. We then used DESeq2 (v1.40.2) (Love et al., 2014) to perform pseudo-bulk DE analysis with a likelihood-ratio test (LRT), where the full model included batch + cluster + lineage, and the reduced model included batch + cluster, in order to detect genes whose expression can be explained by lineage. We used the LRT test by computing pairwise comparisons, contrasting all lineages against each other (CD4vsCD8, CD4vsiNKT, etc.), for each comparison keeping DE genes with an adjusted p-value of 0.01. Then, to extract lineage-specific genes, for each lineage we kept genes that were commonly upregulated in at least 3 or more contrasts. We normalized the raw counts with the rlog function from DESeq2 and batch-corrected them with removeBatchEffect from the limma package (v3.56.2) (Ritchie et al., 2015), before displaying the final list of DE genes on a heatmap with the pheatmap package (v1.0.12).

### Creation of a reference sc-RNAseq mouse T_inn_ dataset

ScRNA-seq data from mouse thymic iNKT, MAIT and γδ T cells (Baranek *et al*., 2020; Chandra *et al*., 2023; Harsha Krovi *et al*., 2020; Koay *et al*., 2019; Lee *et al*., 2020; Legoux *et al*., 2019; Li *et al*., 2022; Maas-Bauer *et al*., 2021; Wang *et al*., 2023) were downloaded from Gene Expression Omnibus repository with the accession numbers GSE137350, GSE172169, GSE179422, GSE130184, GSE152786, the SRA database under the accession code PRJNA549112 or the European Bioinformatics Institute (EMBL-EBI) under the accession number E-MTAB-7704. Data were analyzed using the Seurat package. Analyzed cells were selected to express more than 800 but less than 4200 genes per cell, with less than 5% of mitochondrial reads. Datasets were merged and integrated using the FastMNN algorithm (Haghverdi et al., 2018), using 5000 variable features, k=20 and auto.merge=TRUE. Cell clustering was carried out with a resolution parameter set at 0.5, and potential doublets were detected using scDblFinder (Germain et al., 2021) and subsequently eliminated. To discern differential gene expression between clusters, the FindAllMarkers function was employed, utilizing the MAST algorithm. The analysis considered latent features, specifically the number of genes per cell, and the sample identity, with a log2 fold change threshold of 0.3.

### Interactive data exploration tool

The data from this study is displayed as a ShinyCell application at http://xspeciestcells.chsl.edu. The code for the browser application can be found at https://github.com/meyer-lab-cshl/xspeciestcells-shiny.

### Data availability

Data that support the findings of this study have been deposited in NCBI GEO with the accession code xxx.

### Code availability

Custom analysis code was written in either R (version ≥4.0.3) or python (version ≥3.8). The analysis code is freely available on GitHub: https://github.com/meyer-lab-cshl/xspeciestcells-shiny. The code for the browser application can be found at https://github.com/meyer-lab-cshl/xspeciestcells-shiny.

## Acknowledgments.

We thank members of our laboratories for thoughtful discussions and critical comments on the manuscript; Jennifer Matsuda, James Scott-Browne, Ross Kedl, Jesse Gillis, Douglas Fearon and Leslie Berg for critical comments and support; the Flow Core and the University of Colorado flow cytometry shared resource facility for assistance with cell sorting; the Clinical and Translational Research Centers at the Anchutz Medical campus for collecting blood (supported by NIH/NCATS Colorado CTSA grant UL1 TR002535); the Genomics Core at the University of Colorado Anschutz Medical campus for sequencing and the National Institutes of Health Core facility for CD1d and MR1-tetramers; Rish Prabakar for help in TEC isolation in murine experiments; Sarah Chapin for help in setting up the Shiny app; Eun Seo Park and Jong Kyoung Kim for sharing their mouse γδ T cells seurat object; Daniel P. Caron and Donna L. Farber for sharing their across-tissues T cell AnnData object; Ania Lorenc and Gosia Trynka for sharing their seurat object of human peripheral blood T cells. This work was supported by National Institutes of Health Grants R21AI163454 and R01AI130198 (to L.G.), R56AI155729 (to P.J.N.) and 1R01AI167862-01 (to H.M.). S.C. is supported through the Florence Gould and Annette Kade Fellowships with the Cold Spring Harbor Laboratory of Biological Sciences. This research was further supported by the Simons Center for Quantitative Biology at Cold Spring Harbor Laboratory; the Cold Spring Harbor Laboratory and Northwell Health Affiliation and the Pershing Square Foundation. It was performed with assistance from the Cancer Center Pilot awards Program and the CSHL Shared Resources, including the CSHL Flow Cytometry Shared Resource, which are supported by the Cancer Center support grant 5P30CA045508 as well as assistance from the NIH grant S10OD028632-01 at CSHL. Computational infrastructure was supported by the Alpine HPC system, which is jointly funded by the University of Colorado Boulder, the University of Colorado Anschutz Medical Campus, Colorado State University and the National Science Foundation (award 2201538). Additionally, this study was partly supported by the NIH P30CA046934 Bioinformatics and Biostatistics Shared Resource core at the University of Colorado Anschutz Medical Campus.

## Author contributions

Conceptualization of this study was done by L.L., H.S.K., S.C., H.V.M. and L.G. Analysis of cells and tissues was done by L.L., S.C., J.D., A.S., Y.L. and J.T. Bioinformatics analysis and tool development were carried out by L.G., S.C., H.V.M. and T.B. Funding was acquired by H.V.M. and L.G. Key resources included human tissues from M.S. while some blood CD8 and GD T cells sc-RNAseq data were from W.P and P.N. Supervision of the study was done by H.V.M. and L.G. Writing of the original drafts was done by S.C., H.V.M. and L.G., with review and editing by L.L., S.C., H.S.K., T.B., H.V.M., and L.G.

---

**Supplementary Figure 1: Cell sorting strategy for single-cell sequencing.** Non-myeloid (CD14^−^), non-B-cell (CD19^−^), live cells (viability dye efluor780) from both thymus and blood were sorted into CD4, CD8 and γδ T cells based on CD4^+^CD8^−^, CD8^+^CD4^−^ and CD3^+^TCRγδ^+^ marker expression, respectively. iNKT and MAIT cells were pre-enriched via CD1d-PBS57 and MR1-5OPRU magnetic beads and sorted based on binding to tetramer and TRAV10 or TRAV1-2 stain respectively.

**Supplementary Figure 2: Batch integration and quality control.** A. UMAP projection before and after integration with Harmony, colored by method (RNAseq, RNAseq+VDJseq), donor (1-13), batch (A-I), clusters (1-18) and tissue (thymus and blood). B. Degree of mixing during batch correction and dataset integration measured as the local inverse Simpson’s index (LISI). The top and middle panels show the integration LISI (iLISI), which measures the effective number of datasets within a neighborhood, for thymic and PBMC-derived cells, respectively. Mixing was assessed on batch, donor and method used (as in depicted in A); the lower panel depicts the cell-type LISI (cLISI), to evaluate the accuracy of cell-type assignment. Blue curves indicate LISI before integration, red after integration. C. Cell co-assignment probabilities within (diagonal) and across clusters (off diagonal) assessed by cell bootstrapping and re-cluster. High co-assignment probabilities indicate cluster stability.

**Supplementary Figure 3. Marker gene expression across cell clusters.** A. Reference signature gene expression for clusters in Figure 1C and B. the top five genes that characterize these clusters in this dataset. Top five marker genes for C. iNKT, D. MAIT and E. γδ T cells corresponding to the clusters in Figure 2A, E and I, respectively.

**Supplementary Figure 4. Reproducibility of thymocyte data with human thymus atlas.** A. UMAP representation of our integrated thymocyte data (top) and the Park *et al*. thymocyte data (bottom). Cells are colored by cluster. B. Bubbleplot showing the MetaNeighbor AUROC score for pairwise similarities of our thymocyte clusters with the Park *et al*. (Ref (Park *et al*., 2020)) annotated thymocyte clusters. AUROC scores above 0.9 are written in white text. Marginal bar plots represent the number of cells present in each cluster.

**Supplementary Figure 5: Characteristics of gene expression on integrated dataset.** A. Gene expression projection of signature genes. B. Genes differentially expressed between thymic CD4 and CD8 SP T cells corresponding to clusters c3/c11 and c9/c10 in Figure 1C, respectively.

**Supplementary Figure 6: Projection of GEP12 onto integrated T_inn_ and T_conv_ object.** Each panel shows cells from a given batch, color-coded by the cNMF usage of GEP12. There is a clear separation of batches A-C, E, I and E, F-H, which align with the sequencing method used, RNAseq only or RNAseq+VDJ, respectively (see Supp Table 1).

**Supplementary Figure 7: Gene expression programs (GEP) in thymic T cell types.** Cells are color-coded based on their respective GEP usage (rows) and cell types (columns). GEP usage derived from cNMF usage file.

**Supplementary Figure 8: Effector phenotyping of thymic iNKT and MAIT cells by flow cytometry.** Thymic iNKT (TRAV10^+^ CD1d-PBS57^+^) and MAIT (TRAV1-2^+^ MR1-5OPRU^+^) cells from postnatal thymus were analyzed by flow cytometry for the expression of co-receptors CD4 and CD8; transcription factor PLZF; and effector markers CD161, EOMES, GZMK. iNKT and MAIT cells were pre-enriched via CD1d-PBS57 and MR1-5OPRU magnetic beads.

**Supplementary Figure 9: Effector gene expression programs (GEPs) are consistent across datasets and human tissues.** A. Proportion of genes in each peripheral GEP (3-6) corresponding to genes in public signature gene lists (Poon *et al*. (Poon *et al*., 2023)), Rose et al. (Rose *et al*., 2023)), Cano-Gamez *et al*. (Cano-Gamez *et al*., 2020), Terekhova *et al*. (Terekhova *et al*., 2023) measured by weighted Jaccard Index. For each GEP, the top gene lists with the highest overlap are shown. Tick marks represent the overlap expected from an empirical null distribution (see methods). B. Co-expression of effector GEPs (GEP4-6) and signature gene lists represented on integrated UMAP. For each GEP the co-expression with the gene list corresponding to the highest weighted Jaccard Index (from A) are shown. For the Poon dataset, violin plots on the right represent the effector GEPs scored in cells from the CD4 T_cm/fh_, CD8 MAIT, or CD8 T_em/emra_ clusters, across tissues; the horizontal dashed line is the median score across all clusters and all tissues from the Poon dataset.

**Supplementary Figure 10: Naïve and effector gene and protein expression of adult peripheral blood iNKT cells.** A. Cluster assignment (as in Fig. 4A) and projection of naive-like (GEP3) and effector (GEP4-6) on adult peripheral blood iNKT cells (identified by cell hashtag). B. Gene expression projection of co-receptors (CD4, CD8), transcription factors ZBTB16 (encoding PLZF) and TBX21 (encoding TBET), naïve T cell marker CCR7 and effector markers KLRB1 (encoding CD161), EOMES, and granzymes GZMA, GZMK; C: Flow cytometry of adult peripheral blood iNKT cells (TRAV10^+^ CD1d-PBS57^+^) for a characteristic subset of markers in B.

**Supplementary Figure 11: Naïve and effector gene and protein expression of adult peripheral blood MAIT cells.** A. Cluster assignment (as in Fig. 4A) and projection of naive-like (GEP3) and effector (GEP4-6) on adult peripheral blood MAIT cells (identified by cell hashtag). B. Gene expression projection of co-receptors (CD4, CD8), transcription factors ZBTB16 (encoding PLZF) and TBX21 (encoding TBET), naïve T cell marker CCR7 and effector markers KLRB1 (encoding CD161), EOMES, and granzymes GZMA, GZMK; C: Flow cytometry of adult peripheral blood MAIT cells (TRAV1-2^+^ MR1-5OPRU^+^) for a characteristic subset of markers in B.

**Supplementary Figure 12: Gene and protein expression of adult peripheral blood γδ T cells.** A. Cluster assignment (as in Fig. 4A). B. γ and δ variable segment usage (D-V1-3, G-V9), and C. projection of naive-like (GEP3) and effector (GEP4-6) on adult peripheral blood γδ T cells (identified by cell hashtag). D. Gene expression projection of transcription factors ZBTB16 (encoding PLZF), naïve T cell marker CCR7 and granzymes GZMB, GZMK; E: Flow cytometry of adult peripheral blood γδ T cells. γδ T cells were separated by γ and δ chain usage, either as Vδ2^+^Vγ9^+^, Vδ1^+^, or non-Vδ1^+^ non-Vδ2^+^ cells and analyzed for their expression of the granzymes in D.

**Supplementary Figure 13. Characteristic gene and protein expression of adult peripheral CD4 and CD8 T cells.** A./C. Cluster assignment (as in Fig. 4A) and projection of naive-like (GEP3) and effector (GEP4-6) on adult peripheral blood CD4 and CD8 T cells (identified by cell hashtag), respectively. B./D. Gene expression projection of transcription factors TBX21 (encoding TBET), FOXP3, naïve T cell marker CCR7 and effector marker EOMES, chemokine receptor CXC3CR1 and granzymes GZMA, GZMB, GZMK.

**Supplementary Figure 14. Reference mouse T_inn_ dataset.** A. Integration of single-cell RNAseq data from flow-sorted mouse iNKT, MAIT, or γδ T cells combined from nine independent studies (Refs (Baranek *et al*., 2020; Chandra *et al*., 2023; Harsha Krovi *et al*., 2020; Koay *et al*., 2019; Lee *et al*., 2020; Legoux *et al*., 2019; Li *et al*., 2022; Maas-Bauer *et al*., 2021; Wang *et al*., 2023)) and B. their annotation into 13 clusters, C. spanning across studies and cell lineages. D. Bubble plot of key genes characterizing the 13 clusters.

**Supplementary Figure 15. Gating strategies implemented to identify the various T cell populations for analyses and sorting.** The target (red gate) cell population in each panel is indicated above each panel. iNKT and MAIT cells were pre-enriched by CD1d-PBS57 and MR1-5OPRU tetramers and magnetic beads, respectively.

**Supplementary Figure 16. Determining genes associated with cNMF derived Gene Expression Programs (GEPs).** Gene ranks (sorted most to least associated, x-axis) are displayed against their gene_spectra_score output from the cNMF analysis (y-axis) as black dots. The slope at the first elbow point in the fitted sigmoid curve (red line) was calculated as the minimum threshold for genes to be retained in the a given GEP. The same slope (grey dashed line) was applied to every GEP to prevent bias in ranked gene selection, as the gene ranking between GEPs are not comparable and relative to each GEP.

**Supplementary Table I. Sample overview.** Overview of cell populations collected in this study, their tissue, donor characteristics (Donor, Sex, Age) and analyses methods (Batch, VDJseq).

**Supplementary Table II. A list of genes that are differentially expressed in each of the 18 clusters distributed across both blood and thymus-derived cells.** Cluster-enriched genes by using the FindAllMarkers function in Seurat with test.use = wilcox with log fold change > 0.4 and min.pct = 0.3.

**Supplementary Table III. Gene Signatures used throughout the manuscript.**

**Supplementary Table IV. Ranked gene lists that compose each of the Gene Expression Programs (GEP) determined by cNMF.**

**Supplementary Table V. A list of genes that are differentially expressed in each of the 7 clusters distributed across thymus-derived iNKT cells.** Cluster-enriched genes by using the FindAllMarkers function in Seurat with test.use = wilcox with log fold change > 0.4 and min.pct = 0.3.

**Supplementary Table VI. A list of genes that are differentially expressed in each of the 7 clusters distributed across thymus-derived MAIT cells.** Cluster-enriched genes by using the FindAllMarkers function in Seurat with test.use = wilcox with log fold change > 0.4 and min.pct = 0.3.

**Supplementary Table VII. A list of genes that are differentially expressed in each of the 8 clusters distributed across thymus-derived γδ T cells.** Cluster-enriched genes by using the FindAllMarkers function in Seurat with test.use = wilcox with log fold change > 0.4 and min.pct = 0.3.

**Supplementary Table VIII. A list of genes that are differentially expressed in each of the 4 clusters distributed across blood-derived iNKT T cells.** Cluster-enriched genes by using the FindAllMarkers function in Seurat with test.use = wilcox with log fold change > 0.4 and min.pct = 0.3.

**Supplementary Table IX. A list of genes that are differentially expressed in each of the 4 clusters distributed across blood-derived MAIT cells.** Cluster-enriched genes by using the FindAllMarkers function in Seurat with test.use = wilcox with log fold change > 0.4 and min.pct = 0.3.

**Supplementary Table X. A list of genes that are differentially expressed in each of the 5 clusters distributed across blood-derived γδ T cells.** Cluster-enriched genes by using the FindAllMarkers function in Seurat with test.use = wilcox with log fold change > 0.4 and min.pct = 0.3.

**Supplementary Table XI. A list of genes that are differentially expressed in each of the 6 clusters distributed across blood-derived CD4 T cells.** Cluster-enriched genes by using the FindAllMarkers function in Seurat with test.use = wilcox with log fold change > 0.4 and min.pct = 0.3.

**Supplementary Table XII. A list of genes that are differentially expressed in each of the 6 clusters distributed across blood-derived CD8 T cells.** Cluster-enriched genes by using the FindAllMarkers function in Seurat with test.use = wilcox with log fold change > 0.4 and min.pct = 0.3.

**Supplementary Table XIII. A list of genes that are differentially expressed in each of the 13 clusters distributed across thymus-derived mouse T_inn_ cells.** Cluster-enriched genes by using the FindAllMarkers function in Seurat with test.use = MAST with latent.vars = “orig.ident” and log fold change > 0.3.

